# Replication origin mapping in the malaria parasite *Plasmodium falciparum*

**DOI:** 10.1101/2022.07.27.501677

**Authors:** Francis Isidore Garcia Totañes, Jonas Gockel, Sarah E. Chapman, Richárd Bártfai, Michael A. Boemo, Catherine J. Merrick

## Abstract

The malaria parasite *Plasmodium falciparum* replicates via schizogony: a fundamentally unusual type of cell cycle involving asynchronous replication of multiple nuclei within the same cytoplasm. It also has one of the most A/T-biased genomes ever sequenced. Here, we present the first comprehensive study of the specification and activation of DNA replication origins during *Plasmodium* schizogony. Potential replication origins were found to be abundant, with ORC1-binding sites detected every ∼800 bp throughout the genome. They had no motif enrichment, but were biased towards areas of higher G/C content. Origin activation was then measured at single-molecule resolution via new DNAscent technology that measures fork movement by detecting base analogues BrdU and EdU in DNA sequenced on the Oxford nanopore platform (https://github.com/MBoemo/DNAscent). DNAscent-called origins were found to be much less dense than ORC1-binding sites, with origins activated preferentially in areas of low transcriptional activity. Consistently, replication forks moved fastest through the most lowly transcribed genes, suggesting that conflicts between transcription and origin firing inhibit efficient replication, and that *P. falciparum* has evolved its S-phase to minimise such conflicts.

## INTRODUCTION

The malaria parasite *Plasmodium* causes morbidity and mortality in hundreds of millions of people every year. Severe and fatal malaria is principally caused by the species *Plasmodium falciparum*, which is responsible for ∼600,000 deaths per year (World malaria report 2021, 2021). The extent to which these parasites can replicate in human hosts is very clinically relevant because high parasitaemia is one of the strongest predictors of severe malarial disease (Dondorp *et al*., 2005; Wassmer *et al*., 2015). Accordingly, it is important to understand the cellular and molecular biology of *Plasmodium* replication. Furthermore, the process can be directly targeted with anti-malarial drugs: historically successful combinations of anti-folate drugs worked to inhibit *Plasmodium* DNA replication (Hyde, 2005).

Malaria parasites are fundamentally different from most other pathogens in their replicative processes. Most cells – from bacterial to fungal to protozoal pathogens – replicate via canonical binary fission, whereas these early-diverging protozoans replicate primarily via schizogony. This unique process involves asynchronous replication of multiple nuclei within the same cytoplasm, prior to a single cytokinesis event producing many – and not necessarily 2^n^ – daughter cells (Arnot *et al*., 2011). In fact, this is not the only atypical mode of replication in the complex *Plasmodium* lifecycle. During the single phase of sexual reproduction, which occurs in the gut of the vector mosquito, gametes are produced via a radically different, extremely fast process: 3-fold replication of the genome followed by cytokinesis, all in under 15 minutes (Janse *et al*., 1986; Kawamoto *et al*., 1991).

The demands of both schizogony and gametogenesis are unique. In schizogony, within a single shared cytoplasm, a cell must replicate one genome at first and then as many as dozen genomes just a few hours later, with varying degrees of asynchrony. How this is organised is now being studied in detail at the cell-biological level (Klaus *et al*., 2022; McDonald and Merrick, 2022; Simon *et al*., 2021) but at the molecular level much remains unknown. Most fundamentally, we were interested to know how replication origins are specified and controlled in a genome undergoing several rounds of asynchronous replication and repeated karyokinesis. Our earlier work, using DNA fibre labelling, has shown that replication origin spacing changes across the course of schizogony, with closer-spaced origins occurring in later rounds of replication (Stanojcic *et al*., 2017). Secondly, it remains unknown how replication origins are specified and controlled in the very different situation of gametogenesis, when genome replication is achieved 10-fold more quickly. Our preliminary evidence suggests that many more replication origins must be used at this stage, making origin specification and usage extremely flexible (Matthews *et al*., 2022). Thirdly, some malaria parasites including *P. falciparum* have very unusual genome compositions at over 80% A/T (Gardner *et al*., 2002), the most biased genomes ever sequenced. Other *Plasmodium* species, with similar lifecycles and cell biology, have genomes of only ∼60% A/T (Pain *et al*., 2008). We recently made a preliminary study, again via DNA fibre labelling, of whether origins are specified and controlled similarly in genomes whose nucleotide composition varies so markedly. This suggested that origin spacing is identical during schizogony in A/T-biased versus A/T-balanced species (McDonald and Merrick, 2022).

Replication origin specification has been thoroughly studied in model eukaryotes such as *Saccharomyces cerevisiae* (Donovan and Diffley, 1996), where a complex of proteins based on the origin-of-replication (ORC) complex binds to DNA at specific sites called autonomous replication sequences (ARS). Once these origins are ‘licenced’, they can initiate DNA replication during S-phase (Fragkos *et al*., 2015). In other model eukaryotes including *Schizosaccharomyces pombe, Drosophila* and *Xenopus*, the replication machinery is broadly conserved but ARSs are not. Research continues into the many factors that may specify an origin in these species, from DNA sequence to DNA secondary structure to chromatin architecture (Prioleau and MacAlpine, 2016). In *S. pombe*, for example, A/T-rich sequences are favoured (Dai *et al*., 2005), while in human cells, G-rich G-quadruplex-forming sequences are implicated (Valton and Prioleau, 2016).

In *Plasmodium*, the replication machinery is again broadly conserved, albeit with some protein components missing or unidentified (reviewed in (Matthews *et al*., 2018)). There is limited functional evidence for an ARS, although one computational study has suggested that it may exist (Agarwal *et al*., 2017). ORC subunit 1 (ORC1) plays an important role in DNA replication and has been identified in *P. falciparum* alongside several other ORC subunits (Aurrecoechea *et al*., 2009). It is known to interact with proliferating cell nuclear antigen and both have been shown to co-localise during DNA replication (Gupta *et al*., 2009) suggesting its conserved function in *Plasmodium* as well. In mammals, ORC1 is strongly bound to DNA prior to replication and eventually loses its affinity, or in some cases is degraded, once replication ensues (reviewed in (DePamphilis *et al*., 2006)) making this subunit ideal for genome-wide mapping of potential replication origins (Dellino *et al*., 2013).

In this study, we mapped origins of replication empirically in *P. falciparum* for the first time. In model systems this has been achieved in various ways, including chromatin immunoprecipitation (ChIP) for proteins such as ORC1, and mapping of nascent DNA synthesis by diverse means, including incorporation of modified nucleotides followed by their purification and mapping (Eaton *et al*., 2011). We have combined ORC1-ChIP with a novel application of the recently published nanopore methods (Boemo, 2021; Georgieva et al., 2020; Hennion et al., 2020; Muller et al., 2019; Theulot et al., 2022), which exploits Oxford nanopore long-read sequencing to simultaneously detect and map modified nucleotides in nascent DNA strands. For the first time in any system, we have developed this into a powerful two-nucleotide protocol with new DNAscent software that distinguishes sequential pulses of the thymidine analogues bromo-deoxyuridine (BrdU) or ethyl-deoxyuridine (EdU) in single molecules. When combined with ORC1-ChIP to map all potential origin sites in the *P. falciparum* genome, this provides high resolution and critical insight into replication origin usage in this unusual eukaryote. It also lays vital groundwork for future studies comparing origin usage in different *Plasmodium* species and in different stages of the lifecycle.

## RESULTS

### Distribution of ORC1 is independent of active DNA replication during S-phase in *P. falciparum*

To examine the localisation and intensity of the origin recognition complex subunit 1 (ORC1) in relation to DNA replication in *P. falciparum*, we first C-terminally tagged the *orc1* gene with 3xHA, then transfected a thymidine kinase-expressing plasmid. Thymidine kinase allows the parasites to salvage thymidine analogues such as BrdU or EdU without affecting parasite cell-cycle and survival in the short-term (Merrick 2015). Highly synchronised parasites were generated by purifying late stage schizonts and allowing them to reinvade within a 1-hour window prior to purifying the resultant early-stage parasites. Every 4 hours, from 18 hpi to 46 hpi, an aliquot of the synchronised culture was labelled with BrdU for 30 minutes, then fixed and processed for microscopy. DAPI was used to detect the parasite nuclei, while anti-BrdU and anti-HA were used to detect newly replicated DNA and ORC1, respectively.

Prior to DNA replication and the formation of the first daughter nucleus, ORC1 appeared to be distributed throughout the whole cell, and not strictly localised in the parasite nucleus (Figure 1, 18 hpi). As the ring stage progressed (22 & 26 hpi), it was found generally in the nuclear periphery, consistent with what was reported by Mancio-Silva *et al*. (Mancio-Silva *et al*., 2008). As soon as replication occurred (30 hpi), ORC1 was in closer proximity with the nucleus, and eventually formed distinct spots on the nucleus and its periphery towards the latter part of schizogony (42 hpi onwards). ORC1 signal did not seem to be present in greater amounts in actively replicating nuclei, marked by BrdU (Figure 1, see cell at 34 hpi in which neither of the 2 nuclei has replicated in the prior 30 minutes but ORC1 remains clearly nuclear, and cells at 38 hpi in which a minority of nuclei are replicating but ORC1 is equally present in all nuclei). Quantitative analysis of the signal intensity of BrdU and ORC1 (n = 56-62 cells per timepoint), showed slight fluctuations in ORC1 signal from 18 and 34 hpi, followed by a sharp increase reaching its peak at 46 hpi (Figure 2a). These changes in ORC1 intensity coincided with changes in DNA replication as shown by increases in BrdU and DAPI signal (Figure 2b and 2c). The temporal pattern of ORC1 intensity followed *orc1* expression pattern at the RNA level, with a delay of several hours. The lowest transcription, observed at 20 hpi, and the peak between 35 and 40 hpi (Toenhake *et al*., 2018), were followed by ORC1 intensity at the protein level (Figure 2d and 2e).

**Figure 1:**
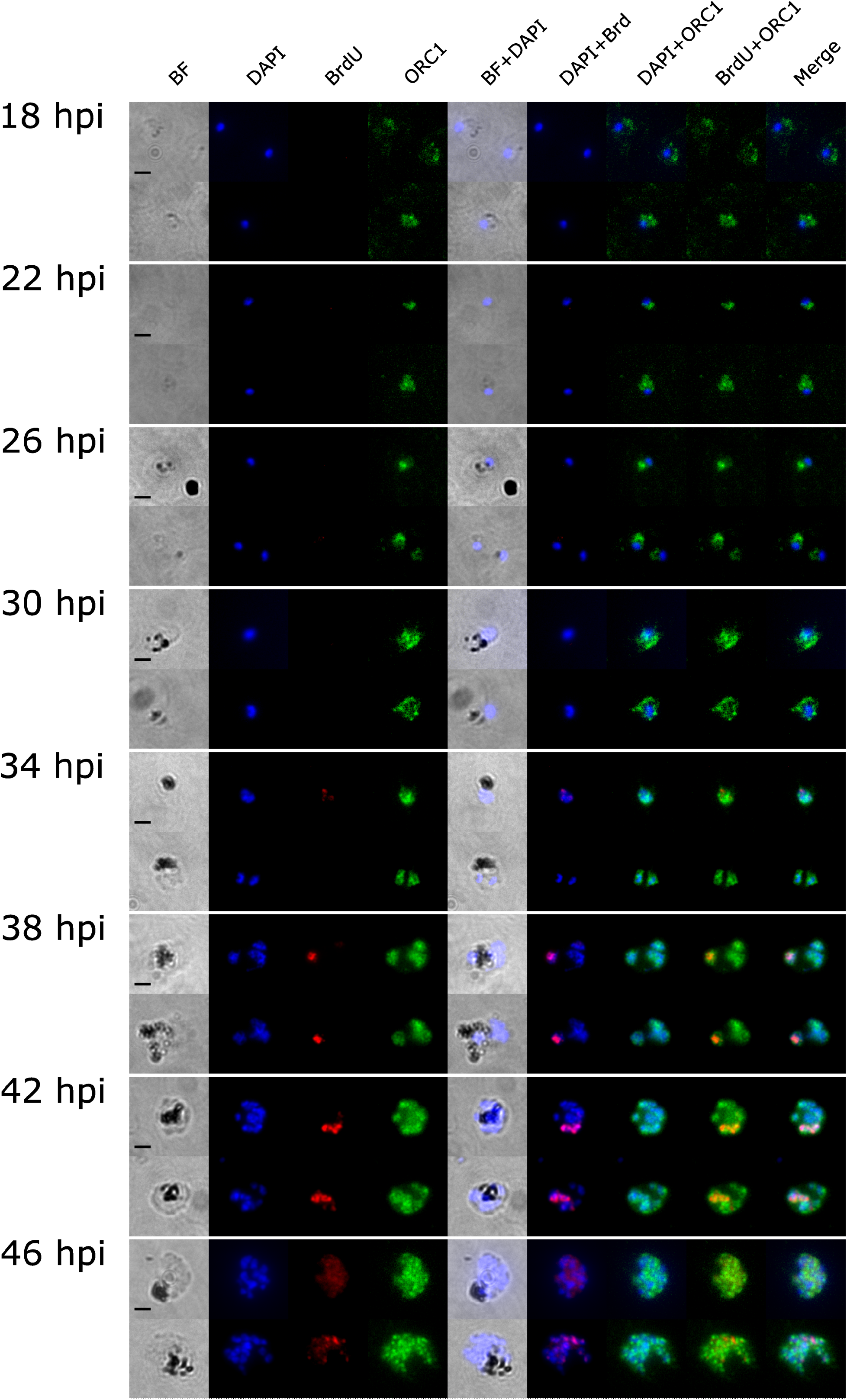
Representative immunofluorescence images showing ORC1 distribution to be independent of BrdU labelling in *P. falciparum* nuclei. Synchronised *P. falciparum* 3D7 ORC1-3xHA + pTK-BSD was incubated in BrdU for 30 mins at 4-hour intervals from 18 to 46 hpi. ORC1 (in green) and BrdU (in red) were probed using anti-HA and anti-BrdU antibodies, respectively. DNA was stained with DAPI (blue). Scale bar (2 μm) applies to all images.

**Figure 2:**
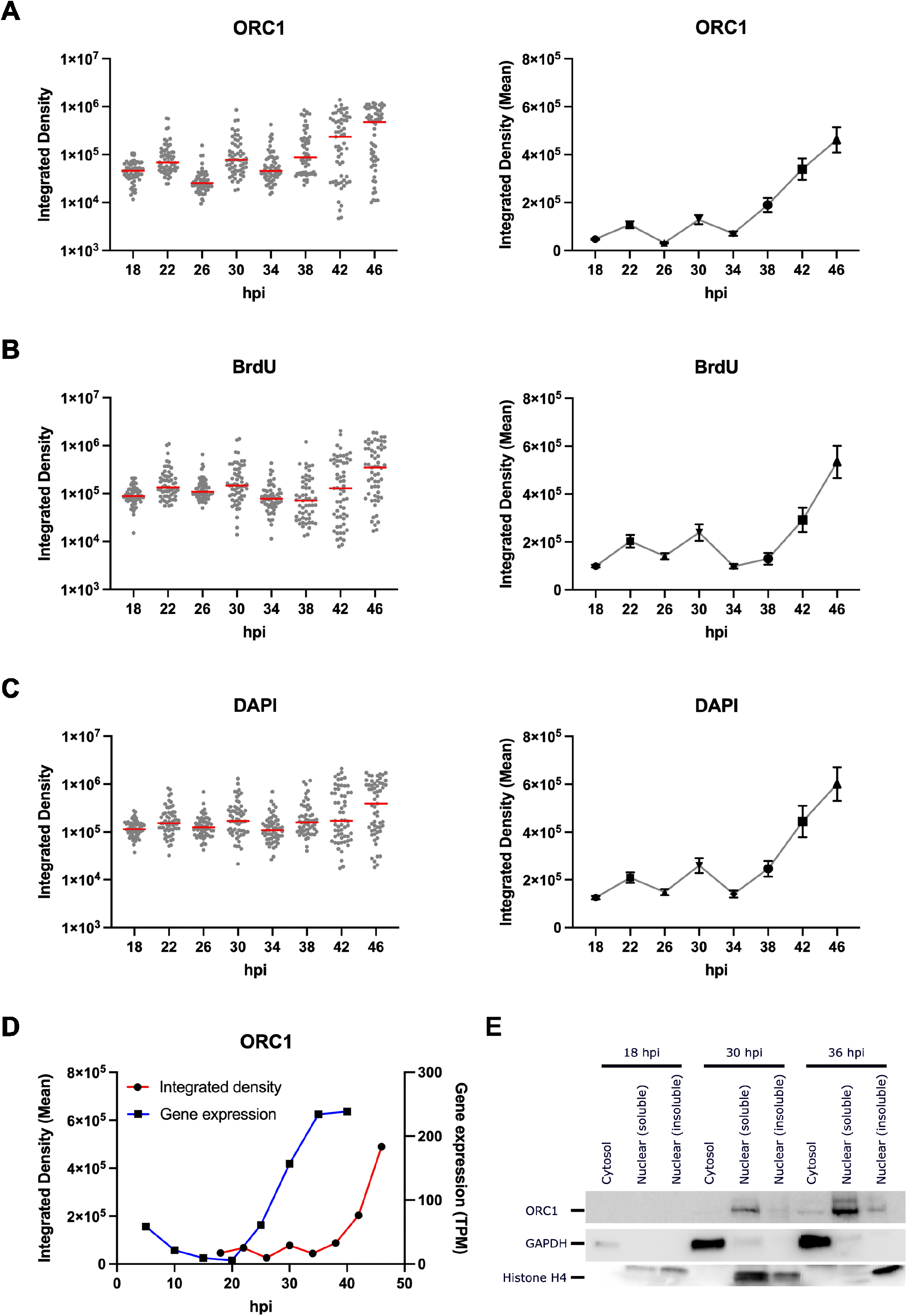
ORC1 abundance increases throughout S phase and follows changes in its transcript abundance. Integrated densities of A) ORC1, B) BrdU, and C) DAPI in tightly synchronised *P. falciparum* 3D7 ORC1-3xHA + pTK-BSD from 18 to 46 hours post invasion (hpi). The median values are represented by red bars in the dot plots (left panel) and line graphs represent mean values ± standard error (right panel) (n = 56-62 cells per timepoint). D) Median ORC1 integrated density in comparison with *orc1* gene expression in transcripts per million (TPM) (Toenhake *et al*., 2018). E) Western blot from fractionated crude protein lysates from synchronised *P. falciparum* 3D7 ORC1-3xHA + pTK-BSD parasites showing ORC1, GAPDH and Histone H4 at 18, 30 and 36 hpi.

### ORC1 chromatin immunoprecipitation reveals that potential origins are densely distributed throughout the genome

ORC1, the largest subunit in the origin recognition complex, plays an important role in DNA replication and has been identified in *P. falciparum* alongside several other ORC subunits (Aurrecoechea *et al*., 2009; Mehra *et al*., 2005). ORC1 binds to replication initiation sites on the genome in preparation for S-phase (DePamphilis *et al*., 2006). Therefore, to identify potential replication initiation sites, we analysed the genomic placement of ORC1 through chromatin immunoprecipitation-sequencing (ChIP-seq) on highly synchronised cultures at two different pre-S-phase timepoints (i.e. 24 and 30 hpi, Figure 3a and b). Data from the two timepoints showed near identical results, with a Spearman R correlation between input-normalised data of 0.924 (Figure 3c). There was a strikingly high density of ORC1 summits, ∼1/kb (median inter-summit distance of 862.5 bp, Supplementary Table 1), with 8,843 peaks identified at 24 hpi and 6,785 at 30 hpi. ORC1 peaks showed a preference for relatively high G/C sequences (Figure 3a and b, median G/C content of 27.0% and 28.4% at 24 and 30 hpi, respectively). ORC1 was enriched in, but not limited to, the coding sequences which are likewise higher in G/C than the total genome (Figure 3d). Furthermore, we observed ORC1 enrichment in sub-telomeric regions, consistent with previously published data (Deshmukh *et al*., 2012; Mancio-Silva *et al*., 2008) (Figure 3a).

**Figure 3:**
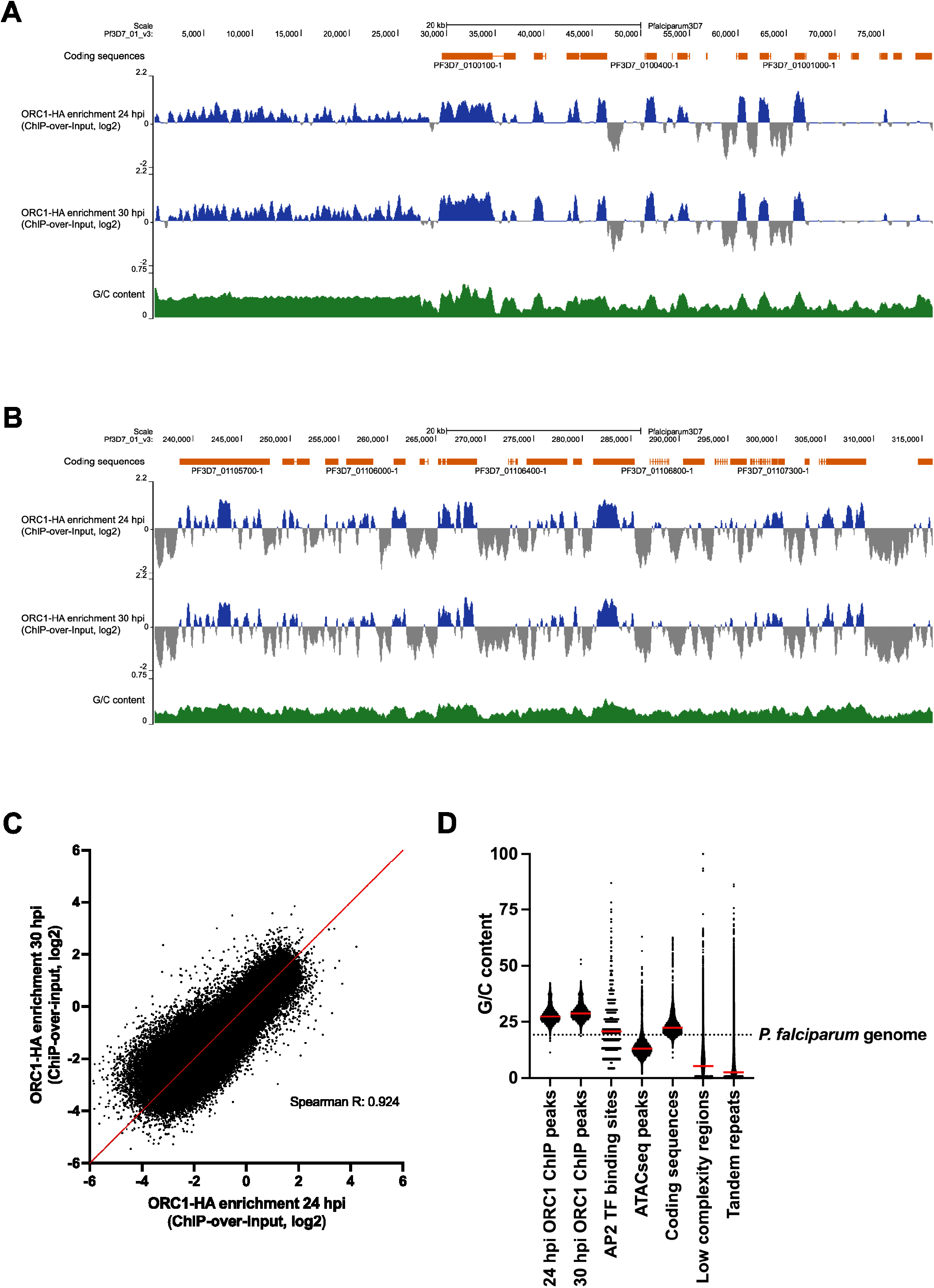
ORC1 ChIP-seq shows consistent ORC1 localisation in relatively high G/C regions. A) ORC1 ChIP log2ratios (normalised against input) at 24 and 30 hpi (middle two panels, in blue) on the first 80 kb of chromosome 1 are visualised here alongside coding sequences (top panel, in orange) and G/C content (bottom panel, in green). B) ORC1 ChIP log2ratios at 24 and 30 hpi in a non-telomeric region. C) Scatterplot where points are the average log2ratios of ORC1 ChIP data over each non-overlapping 100 bp window in the genome at 24 hpi (x-axis) and 30 hpi (y-axis). The line where x=y is show in red. D) Comparison between the G/C content at ORC1 ChIP-seq peaks and the G/C content at ATAC-seq peaks, predicted binding sites of AP2 transcription factors, coding sequences, low complexity regions, and tandem repeats. The overall G/C content of *P. falciparum* is represented by the dashed line while red bars represent the median G/C content.

**Table 1:**
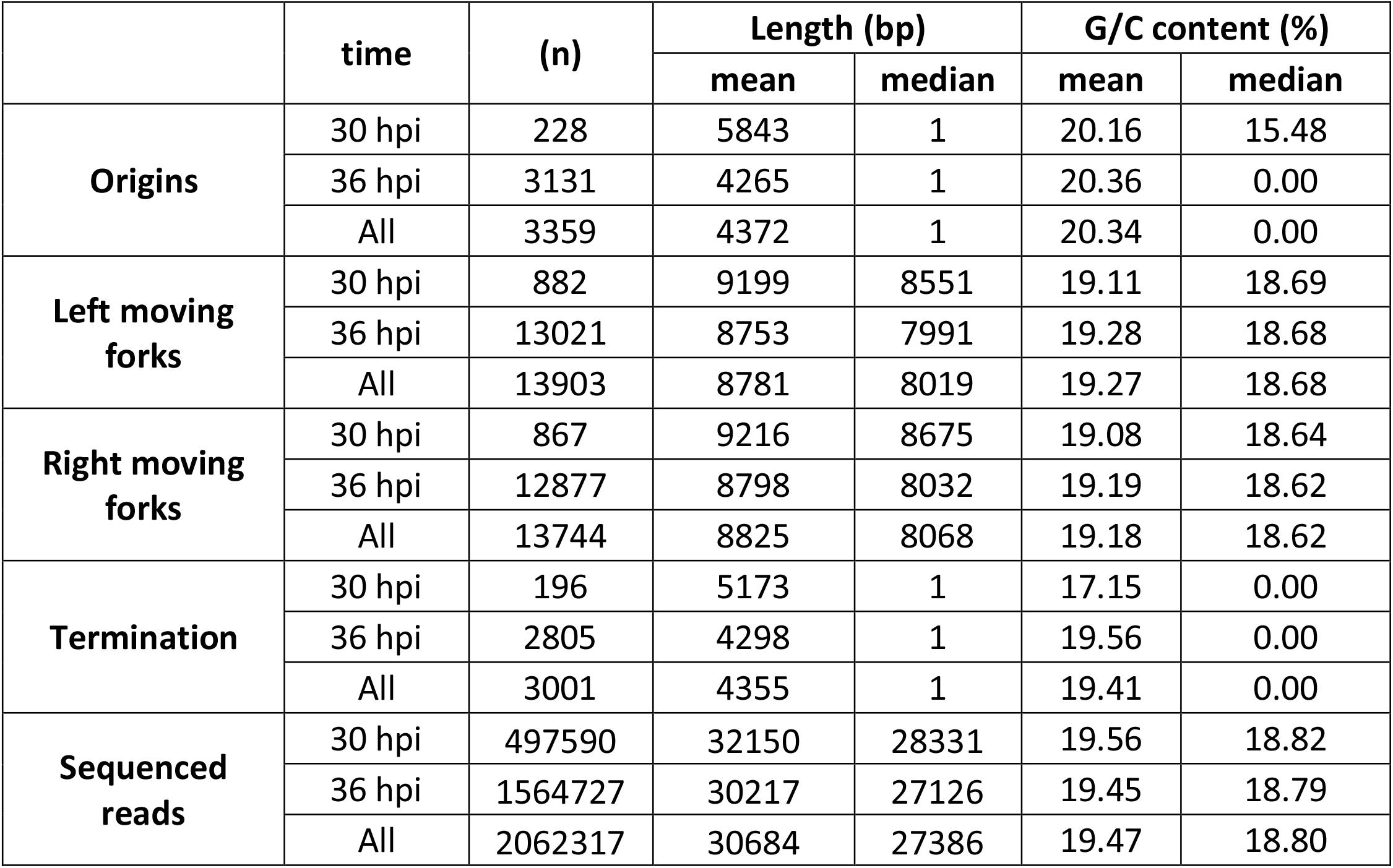
DNAscent identifies origins, ongoing replication forks and terminations. Total number of origins, left-/right-moving forks, termination sites identified by DNAscent v3.0.2, and the total quality sequenced reads with the corresponding mean and median lengths and G/C content. Note that origins include both those that are mapped to a centre point (1 bp) and those derived from two separated left- and right-moving forks, in which case the origin is not mapped to a point, but to the entire inter-fork region.

ORC deposition/binding is not sequence-specific in most organisms, but the *S. cerevisiae* autonomously replicating sequence (ARS) is an exception, and similar sequences have been proposed to act as ORC landing pads in *P. falciparum* (Agarwal *et al*., 2017). We therefore conducted a *de novo* motif search on ORC1 ChIP sequences ± 50 bp from the summits using HOMER motif analysis software. We identified 12 unique motifs from the sequences that bound to ORC1 at 24 and 30 hpi (Supplementary Table 2). The overall mean weighted G/C content of these motifs was 28.7%, with a much higher G/C content for motifs found at 24 hpi (30.7%) compared with those at 30 hpi (25.1%). These findings are not consistent with previously reported sequences such as the *S. cerevisiae* autonomously replicating sequence (ARS) (Dhar *et al*., 2012), or the proposed variant ‘PfARS’ (Agarwal *et al*., 2017) that are less G/C rich (0 to 18.2% G/C). Furthermore, motifs identified were only moderately enriched and are therefore unlikely to serve a similar function as ARS.

In contrast to the G/C-poor ARS, G-rich sequences that can form G-quadruplexes (G4s) have been proposed to promote ORC binding in human cells (Hoshina *et al*., 2013; Prorok *et al*., 2019). Therefore, we also calculated the enrichment of G4-forming motifs in ORC1 ChIP peaks using G4 Hunter (Bedrat *et al*., 2016; Brazda *et al*., 2019). Compared with the overall occurrence of G4-forming motifs in the whole *P. falciparum* 3D7 genome, there was a small enrichment of 2.05 and 2.82-fold in ORC1 ChIP peak sequences at 24 and 30 hpi, respectively (Supplementary Table 3).

Finally, the chromatin landscape has been proposed to influence ORC distribution in some systems (Eaton *et al*., 2011), so we compared ORC1’s whole genome distribution with previously published epigenetic landscapes such as variant and modified histone placements, chromatin accessibility and predicted binding sites of the apicomplexan apetala2 (ApiAP2) transcription factors (Campbell *et al*., 2010). We found no correlation or a very weak negative correlation with most histone and modified-histone distributions (Spearman R: -0.153 to 0.033). Interestingly, we observed more negative correlation between ORC1 placement and predicted AP2 binding sites (Spearman R: -0.208 to -0.192) and a moderate-to-strong negative correlation with accessible chromatin (Spearman R: -0.630 to -0.283) (Supplementary Table 4). Open chromatin identified through the Assay for Transposase-Accessible Chromatin (ATAC-seq) has been shown to correlate with transcription factor binding (Toenhake *et al*., 2018). This suggests a potential inverse relationship between replication origin placement and transcription dynamics. However, since ORC1 was generally associated with areas of high-G/C sequence and with coding sequences, the G/C content at the associated genomic locations could be a potential confounding factor (median G/C at ATAC-seq peaks: 13.1%).

We also compared ORC1 distribution with heterochromatin protein 1 (HP1) since these proteins have been demonstrated to co-localise or interact with each other in *Plasmodium* and other organisms (Deshmukh *et al*., 2012; Pak *et al*., 1997; Prasanth *et al*., 2004; Shareef *et al*., 2001). Consistently, we observed weak to moderate positive correlation between ORC1 and HP1 (Spearman R: 0.3402 to 0.5737). This positive correlation could be related to the fact that ORC1 and HP1 are associated with telomeres (Hoeijmakers *et al*., 2012; Mancio-Silva *et al*., 2008); however, we also calculated correlation between ORC1 and HP1 outside the telomeric regions and still found positive correlation (Spearman R: 0.2555 to 0.5095) (Supplementary Table 4).

### Detection of replication origin activity with single-molecule resolution

Our ORC1 ChIP-seq data identified a high density of ORC1 placement throughout the genome (∼1 per kb). This is a population-level experiment, so every site may not be occupied in every parasite, but nevertheless it was notable that our previous data showed a lower density of active origins on actively replicating DNA fibres (∼1 per 65kb) (Stanojcic *et al*., 2017). This difference in the density of ORC1 placement and active origin firing suggests that parasites license a large excess of origins and fire only a small subset, similar to other organisms (Bell and Labib, 2016; Ge *et al*., 2007). To verify this, we utilised long-read Oxford nanopore sequencing of genomic DNA to accurately identify active replication with sequence-specificity and single-molecule resolution. Tightly synchronised thymidine kinase-expressing *P. falciparum* parasites at early and mid S-phase were sequentially pulse-labelled with two different thymidine analogues, EdU and BrdU, for a total of 15 minutes followed by high-molecular-weight DNA preparation and sequencing. The thymidine analogues would thus mark DNA replication forks that had been active within the past 15 minutes, allowing us to confidently identify replication fork directionality and speed, and to infer the placement of active replication origins (Figure 4a and b). Sequencing data was analysed using DNAscent software v3.0.2, an overhauled version of its predecessor, which originally detected BrdU nucleotide incorporation into replicating DNA and identified fork direction and speed via the changing gradient of BrdU incorporation in the nascent strand (Boemo, 2021). To develop the most recent version, DNAscent was trained to distinguish between EdU and BrdU on the same molecule in order to more accurately and reliably identify fork directionality and origins.

**Figure 4:**
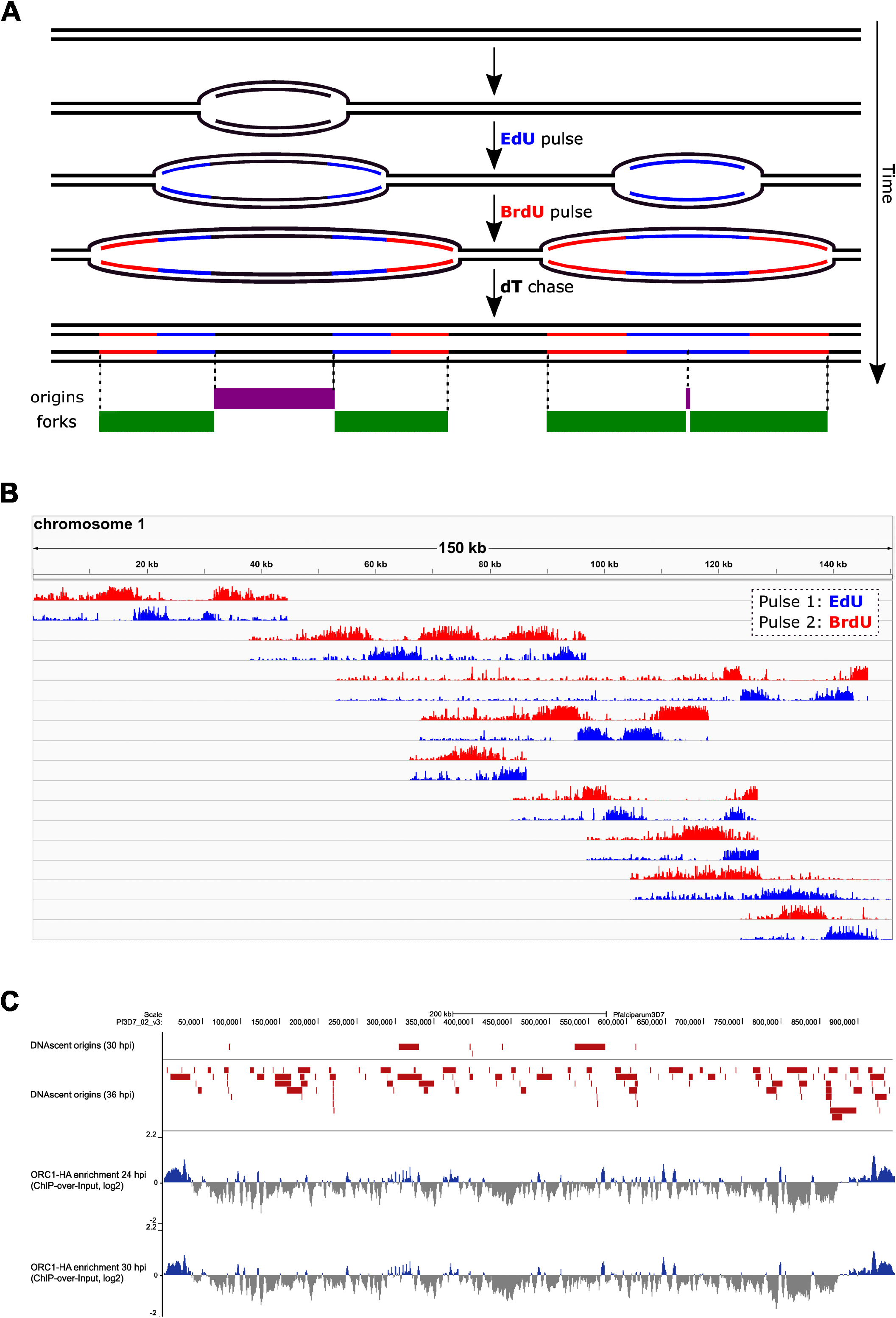
DNAscent v3.0.2 identifies actively replicating DNA and replication origins. A) Schematic of the pulse-labelling protocol used to prepare sequencing reads for analysis by DNAscent v3.0.2. The thymidine analogue EdU (blue) is pulsed into replicating DNA where it is incorporated into the nascent strand by replication forks. This is followed by the pulse of a second thymidine analogue BrdU (red) to indicate fork direction (as in established DNA fibre-labelling protocols), which is in turn followed by a thymidine chase. While this study used an EdU pulse followed by a BrdU pulse, the order is interchangeable and the DNAscent v3.0.2 software can analyse reads with analogues pulsed in either order. Coordinates of origins and forks called by the DNAscent software are represented in purple and green, respectively. B) Nine single molecules sequenced on the Oxford nanopore MinION platform that mapped to *P. falciparum* chromosome 1 and were analysed with DNAscent v3.02. Similar to the visualization in (Boemo, 2021), each molecule is represented as two bedgraph tracks showing the probability of BrdU (red) and EdU (blue) called by DNAscent v3.0.2 at each thymidine position in the read. C) Visualisation showing fired origins called by DNAscent at 30 and 36 hpi on chromosome 2 (red) with ORC1 ChIP peaks (positive peaks in blue, negative peaks in grey) for comparison.

### Only a subset of potential origins are used as active replication origins during *P. falciparum* schizogony

We conducted a series of nanopore sequencing experiments and generated more than 2 million quality reads, and more than 63 billion bases sequenced, i.e. ∼2,700 times genome coverage. In contrast to ChIP-seq where aligned sequences and the corresponding sequence enrichments (i.e. peaks and summits) represent population level data, analysis of these nanopore sequencing reads with DNAscent v3.0.2 reveals snapshots of DNA replication dynamics on single molecules.

Table 1 summarises the total number of left- and right-moving forks, origins and replication terminations detected by DNAscent. About two-thirds (2151 out of 3359) of the origins identified fired during the EdU pulse (Figure 4a, right replication schematic), while the rest fired before the pulse (Figure 4a, left replication schematic). Origins that fired during the EdU pulse were assigned a 1 bp coordinate, representing the centre of the EdU-labelled region, while origins that fired before the pulse – and may therefore have been anywhere between the two EdU-labelled regions – are represented by the entire region between the left- and right-moving forks. Mean G/C content in these identified origins was slightly higher (20.3%) compared to left- and right-moving forks (19.1 to 19.3%) which have G/C content relatively similar to that of the whole genome (19.3%). We detected significantly fewer fired origins at 30 hpi and 36 hpi compared with the total number of ORC1 ChIP peaks. Importantly, neither set of sites is necessarily fully and identically present in every single cell, since both these techniques provide whole-population measurements (however, where adjacent replication forks and origins appear on the same nanopore sequencing read, they must come from a single nucleus).

This confirms that although ORC1 is widely and consistently distributed throughout the genome prior to and at the beginning of S-phase, only a few of the origins are active at mid-S-phase and even fewer at the beginning of S-phase. We identified a total of 228 origins at 30 hpi and 3,131 at 36 hpi, which (after adjusting for the total number of sequenced reads) is over 4 times more at mid S-phase than at early S-phase, suggesting that more potential origins are competent to fire at mid-S-phase (Table 1). Furthermore, there was clear evidence by mid S-phase for ‘efficient’ versus ‘inefficient’ origins, with some regions being detected as an active origin on many separate sequence reads, and others only once (Figure 4c).

### Overlap of active replication origins with ORC1 binding sites and other genomic features

We did not see any significant correlation between active origins and ORC1 ChIP sites – perhaps because active origins were 1-2 orders of magnitude less abundant than ORC1 binding sites. There was also no correlation between active origins and histone markers, predicted AP2 transcription factor binding sites, and chromatin accessibility. However, there was negative correlation between active origins and gene expression, as measured by previous transcriptomic studies conducted across the lifecycle (correlation coefficients of -0.1515 and -0.1736 at 30 and 36 hpi (Supplementary Table 5)). To examine transcriptional activity at these active origins, we calculated the median expression of genes overlapping with origins that fired at 30 and 36 hpi and compared those against the median gene expression of all genes at the closest available time point in published data, i.e., 30 and 35 hpi (Toenhake *et al*., 2018). Median gene expression was significantly lower at active origin sites during both early and mid-S-phase (p-values: 9.88 e-14 and 8.27 e-43, respectively, Figure 5a). This suggests that origins are more likely to fire when positioned in genes that are low-expressing, and that this preference is even greater in origins fired early in schizogony. This is in stark contrast with what has been observed in human cells, where origins located in highly expressed genes tend to fire early during S-phase (Dellino *et al*., 2013).

**Figure 5:**
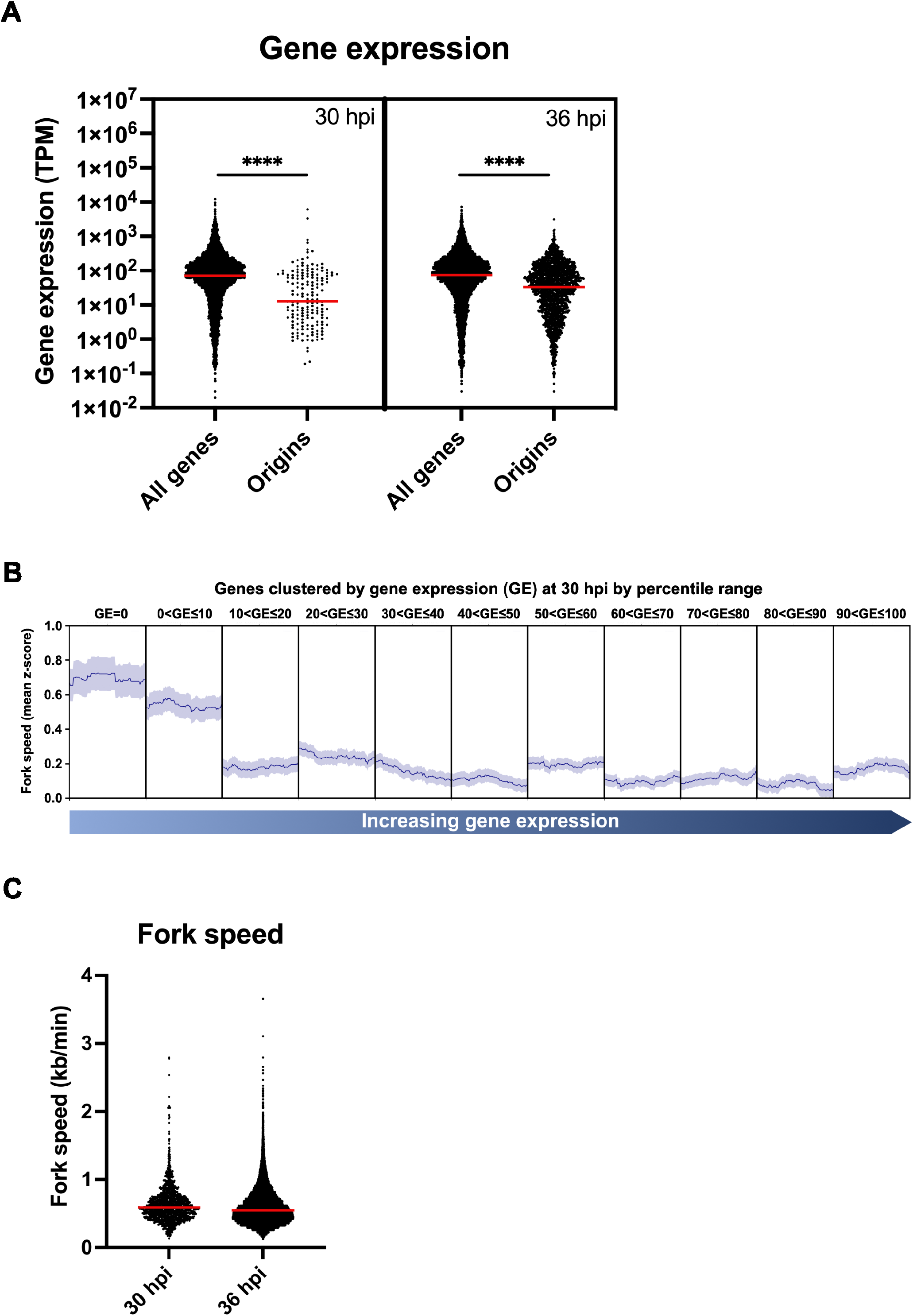
Median gene expression is lower in active origins, while fork speed is faster in lowly transcribed regions. A) Gene expression (in transcripts per million, TPM) of genes that overlapped with active origins (± 50 bp from the origin centre) at 30 and 36 hpi alongside expression of all genes at those time points. Red bars represent median gene expression. B) Mean fork speed z-score (dark blue line), representing the average deviation of the raw fork speed from the overall mean fork speed, mapped along coding sequences of genes clustered by gene expression at 30 hpi. The coding regions of genes within each cluster were scaled, with the left and right borders of the plot corresponding to the transcription start site and transcription end site, respectively. The standard error of the mean is shown as light blue shading above and below the mean fork speed. C) Dot plot showing fork speed at 30 and 36 hpi, with the median represented by the red bar (median at 30 hpi: 0.589 kb/min, at 36 hpi: 0.548 kb/min).

GO term analysis of genes at the active origins revealed 4 out of the top 5 curated GO processes to be related to the virulence gene family *var*, which is known to be largely transcriptionally silent (Supplementary Table 6). ORC1 has been suggested to have an important role in *var* gene regulation (Deshmukh *et al*., 2015). Interestingly there was a positive correlation between ORC1 ChIP and origin activity at *var* genes (0.1298 to 0.3065), which differs from the lack of correlation across the whole genome (-0.0733 to -0.0014) (Supplementary Table 5).

Finally, building on the finding that origins fire preferentially in genes with low expression, we checked whether the progress of replication forks might be affected by transcription-replication conflicts. Indeed, we found that the speed of forks was slower as they traversed the most highly-expressed genes and faster in the lowest-expressed as well as transcriptionally silent genes at 30 hpi (Figure 5b). Median fork speed was also significantly faster at the beginning than the middle of schizogony: 0.589 kb/min at 30 hpi versus 0.548 kb/min at 36 hpi (p-value = 3.24 e-08) (Figure 5c). The same trend was reported in previously published data from DNA fibre combing, although the absolute speeds measured by that method were markedly faster (1.388 kb/min at early trophozoite stage and 1.176 kb/min at mid/late trophozoite stage) (Stanojcic *et al*., 2017). However, the fork speeds calculated here are much closer to those calculated in a more recent publication that utilised DNA spreading rather than combing (0.6-0.7 kb/min) (McDonald and Merrick, 2022).

### Modelling of replication origin activity in *P. falciparum*

To check whether the single-molecule replication parameters measured here were consistent with the observed time periods taken to replicate a whole nucleus (Klaus *et al*., 2022; McDonald and Merrick, 2022), we created a simple stochastic model of *P. falciparum* DNA replication in the Beacon Calculus (Supplementary Figure 1) (Boemo *et al*., 2020). For simplicity, we used only *P. falciparum* chromosome 1, taking the median spacing between licensed origins on chromosome 1 as that measured by ORC1 ChIP (700 bp) and the average replication fork speed as that measured by DNAscent at 30 hpi (0.64 kb/min). Despite being deliberately simple, the model (Figure 6) showed remarkably good agreement with the distribution of inter-origin distances and fork speeds measured by fibre spreading, as well as the total replication time per nucleus of 40-75 minutes, measured by McDonald and Merrick (McDonald and Merrick, 2022).

**Figure 6:**
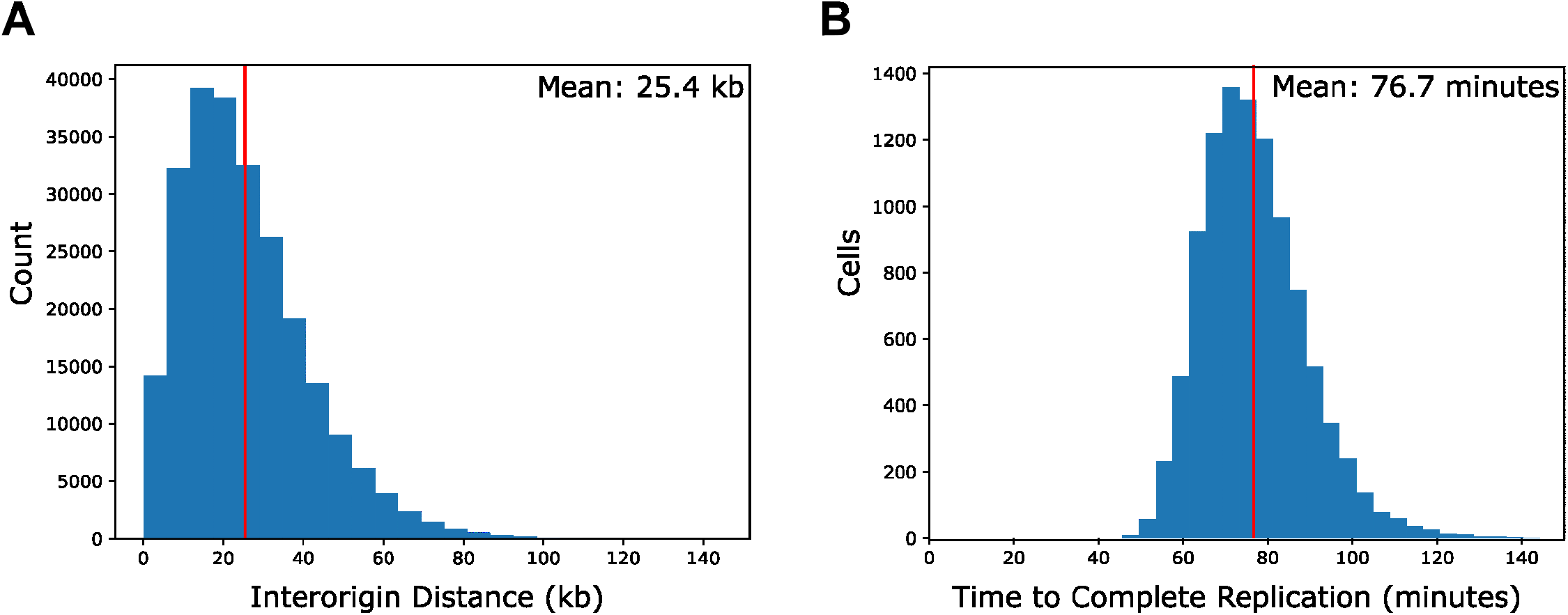
Beacon calculus model showed good agreement with the distribution of inter-origin distances and fork speeds in DNA fibre-spreading data, as well as with the total replication time per nucleus. Distribution of (A) inter-origin distances, and (B) total replication time of non-telomeric regions for *P. falciparum* chromosome 1 found using the Beacon Calculus model shown in (Supplementary Figure 1). The model has a spatial distribution of 100 bp, and distributions are plotted from 10,000 simulations of chromosome 1 DNA replication. Vertical red lines show the mean of each distribution. Simulation of the model was done with the Beacon Calculus Simulator software v1.1.0.

## DISCUSSION

This study represents the first empirical analysis of DNA replication origin activity in the unusual protozoan parasite *Plasmodium*. To achieve this, we combined ChIP-seq mapping of ORC1 binding sites with newly evolved DNAscent software to map active origins on single DNA molecules.

We found that *Plasmodium* ORC1 does not appear to organise the asynchronous replication of multiple nuclei within the same cell that is characteristic of schizogony. ORC1 entered the nucleus before S-phase started and evidently remained there as the number of nuclei increased, being synthesised continuously, presumably to keep pace with the growing number of total genomes per cell. By contrast, proliferating cell nuclear antigen (PCNA), appears to have a limited supply in the parasite cell and shuttles between nuclei, providing a marker for which ones are actively replicating (Klaus *et al*., 2022). There is also no clear evidence in our cell-level data for ORC1 degradation between rounds of replication, as occurs in some systems (although previous data suggest that ORC1 is degraded during the late schizont stage, after S-phase has finished (Deshmukh *et al*., 2015)). In a cell that does not conform to once-and-only-once genome replication, it should be more efficient to keep ORC bound to each genome for the next round until S-phase is completed, presumably while adding newly-synthesised ORCs to each daughter genome. Alternatively, ORC may be evicted into the nucleoplasm at the end of each replicative round, partitioned into daughter nuclei and then re-bound to DNA for the subsequent round. Our ChIP data do not address this because we mapped ORC1 only prior to and at the onset of the first replicative round. In fact, if ORC was loaded onto DNA exclusively at this stage and then progressively evicted after origin activation during each of 4-5 replicative rounds, this might explain why ORC1 binding sites detected by ChIP were extremely abundant. However, the production of additional ORC1 protein throughout S-phase, as more genomes are generated, would then be unnecessary, arguing against this model.

ChIP revealed that ORC binding sites are extremely abundant and not sequence-specific in *P. falciparum*. Nevertheless, the appearance of clear ChIP peaks showed that binding is not totally random: there was a clear preference for G/C enrichment. This is unusual, since ORC in most species – including *S. cerevisiae* and *S. pombe* – prefers A/T-rich sequence (Mechali, 2001). However, even a relative ‘enrichment’ in a genome of only ∼19% G/C remains low, and it is unlikely to signify a preference for G-quadruplex DNA structures, as suggested in human cells (Hoshina *et al*., 2013; Prorok *et al*., 2019). A basic G/C preference could explain why G-quadruplex-forming motifs were somewhat enriched in the ChIP peaks, but the density of peaks was far greater than the predicted density of quadruplex motifs in the *P. falciparum* genome (Gage and Merrick, 2020; Gazanion *et al*., 2020). In fact, a simple G/C binding preference in ORC1 may be sufficient to generate clear sites of enrichment in what is a highly A/T-biased genome.

Surprisingly, no other features of the chromatin landscape seemed to determine ORC1 distribution. In other systems, ORC1 has been associated with intergenic regions (which was not the case here), as well as transcription start sites, acetylated histones, open chromatin, and CpG islands (which are near-absent in *P. falciparum* (Bunnik *et al*., 2013)). ORC1 showed no correlation with epigenetic marks and a negative correlation with transcription factor binding sites and accessible chromatin, although this could be confounded by their opposite bias towards A/T-richness. The exception was a positive correlation between ORC1 and HP1, a marker of heterochromatin: this correlation persisted even outside the telomeres, where ORC1 may play separate, non-S-phase roles in maintaining telomere structure and silencing (Deshmukh *et al*., 2015; Mancio-Silva *et al*., 2008).

ORC1 was evidently bound to the genome very densely at G/C-enriched sites – although it may not be present at every site in every cell since ChIP-seq is a population-level technique. Nevertheless, as in most eukaryotic cells, it seems that only a fraction of these sites were used as active origins. Activation did appear to be determined by the chromatin environment – more precisely, the transcriptional landscape. However, instead of a bias towards firing origins in highly transcribed genes (Gilbert, 2002), *P. falciparum* showed the opposite bias. An imperative to reduce transcription/replication conflicts by preferentially activating origins in low-expressed genes may drive this bias. The molecular mechanism remains unknown, but in a cell that is probably fundamentally replication-stressed (Stanojcic *et al*., 2017) and limited in its ability to activate checkpoints for DNA damage repair (Matthews *et al*., 2018), it could be particularly important to evolve a system that avoids conflict between RNA polymerases and actively firing origins.

ORC1-ChIP yielded so many sites that there was no overall correlation between these sites and active origins identified via DNAscent – this is in contrast to *S. cerevisiae* (Xu *et al*., 2006), where a greater proportion of origin sites are actively used, but more similar to metazoans, where only 5-20% of ORC sites may be used (Hamlin *et al*., 2008). Some sites were used more efficiently than others, similar to the situation in yeast and other systems (Mechali, 2001), because some active origins were detected in similar positions on many DNA strands, whereas others were not. It is not yet clear what determines origin efficiency in *P. falciparum*. Lack of transcription may be entirely responsible, or there may be other factors besides those tested here. Interestingly, the large *var* gene family may present an extreme example of the preference for firing origins in transcriptionally inactive genes. Here, amongst ∼60 genes that are almost entirely silenced, ORC1 placement and origin activation correlated well, whereas in the rest of the genome, which is more variably transcribed, ORC1 sites were so dense versus the density of active origins that there was no overall correlation.

Consistent with the concept that *Plasmodium* has evolved to minimise transcription/replication-origin conflicts, we found that replication forks moved slowly through the most highly transcribed genes, suggesting that such conflicts do inhibit efficient DNA replication. The ability to measure replication fork velocity is another powerful aspect of the DNAscent software, and is orthogonal to the more traditional measurement of fork velocity via DNA combing. Fork velocities in *P. falciparum* DNA were strikingly lower when measured by DNAscent or by DNA fibre spreading (McDonald and Merrick, 2022) than they were by the usual ‘gold standard’ of DNA combing (Stanojcic *et al*., 2017). With most genomes DNA combing stretches the DNA fibres at a reproducible 2 kb/μm (Michalet *et al*., 1997). However, *P. falciparum* DNA must be alkali-treated prior to combing to remove the haemozoin that interferes with combing (Stanojcic *et al*., 2017), and this may result in over-stretching and hence an overestimation of fork velocity. Conversely, both DNA spreading and DNAscent may give an underestimate because they rarely measure very long fibres and hence neglect the longest forks, whereas hundreds of kilobases can often be measured contiguously on combed DNA. In fact, a combination of both these issues could contribute to disparities in measured fork velocity: 1.0-1.4 kb/min as measured by combing, versus 0.6-0.7 kb/min by spreading or DNAscent. Assuming that the lower measurements are correct, then they are strikingly slow compared to either human cells or yeast. High rates of fork asymmetry and stalling were measured in our previous DNA-combing study (Stanojcic *et al*., 2017) (these are independent of absolute velocity), and they suggest that *P. falciparum* operates under considerable replication stress – possibly exacerbated by its A/T-rich and highly repetitive genome, which may cause DNA polymerases to move unusually slowly.

In addition to providing the first map of replication origin activity in *P. falciparum*, this study paves the way to investigate other interesting aspects of malaria parasite biology. Firstly, is origin specification similar in other *Plasmodium* genomes such as *P. knowlesi*, which are not as G/C-poor as *P. falciparum* (Pain *et al*., 2008)? Here, a simple G/C preference in ORC1 binding might not be sufficient to yield clear binding sites. Secondly, how do the demands of gametogenesis – where genome replication is extremely fast but there is little active transcription (Matthews *et al*., 2022) – differ from those of erythrocytic schizogony? Gametogenesis may resemble an extreme version of replication in early *Xenopus* or *Drosophila* embryos, where origins fire every 10-20 kb (Hyrien *et al*., 1995). Indeed ORC1 binding sites may be extremely dense in this system simply because they are needed uniquely in male gametes, whereas they are never all activated in schizogony. It is evident that much remains to be studied about the very unusual cell cycles pursued by malaria parasites.

## ACKNOWLEDGEMENTS

We are grateful to Dr Craig Duffy for early cloning work towards the tagged ORC1 line. This project has received funding from the European Research Council (ERC) under the European Union’s Horizon 2020 research and innovation programme (Grant agreement No. ERC-2016-COG 725126). This project has also received funding from the Isaac Newton Trust (Grant number: 19.39B), Royal Society (Grant number: RGS\R1\201251) and from the Department of Pathology, University of Cambridge. We acknowledge the support from the MSCA ITN Cell2Cell (funded by the European Union’s Horizon 2020 Research and Innovation programme, grant number 860675) for the fellowship of J.G.

## AUTHOR CONTRIBUTIONS

C.J.M. conceptualised the research and conducted overall project administration. F.I.G.T., S.E.C., and M.A.B. were involved in designing the methodology. F.I.G.T., J.G., and M.A.B. conducted the research investigation and data curation. Software development was done by M.A.B. Formal analysis and data visualisation was done by F.I.G.T., J.G., S.E.C. and M.A.B. R.B., M.A.B., and C.J.M. supervised the project. F.I.G.T. and C.J.M. wrote the original draft. F.I.G.T., R.B., M.A.B., and C.J.M. were involved in the review and editing of the final manuscript. R.B., M.A.B. and C.J.M. acquired funding for the project.

## DECLARATIONS OF INTEREST

The authors declare no competing of interest.

## INCLUSION AND DIVERSITY

One or more of the authors of this paper self-identifies as an underrepresented ethnic minority in science.

## STAR METHODS

### KEY RESOURCES TABLE

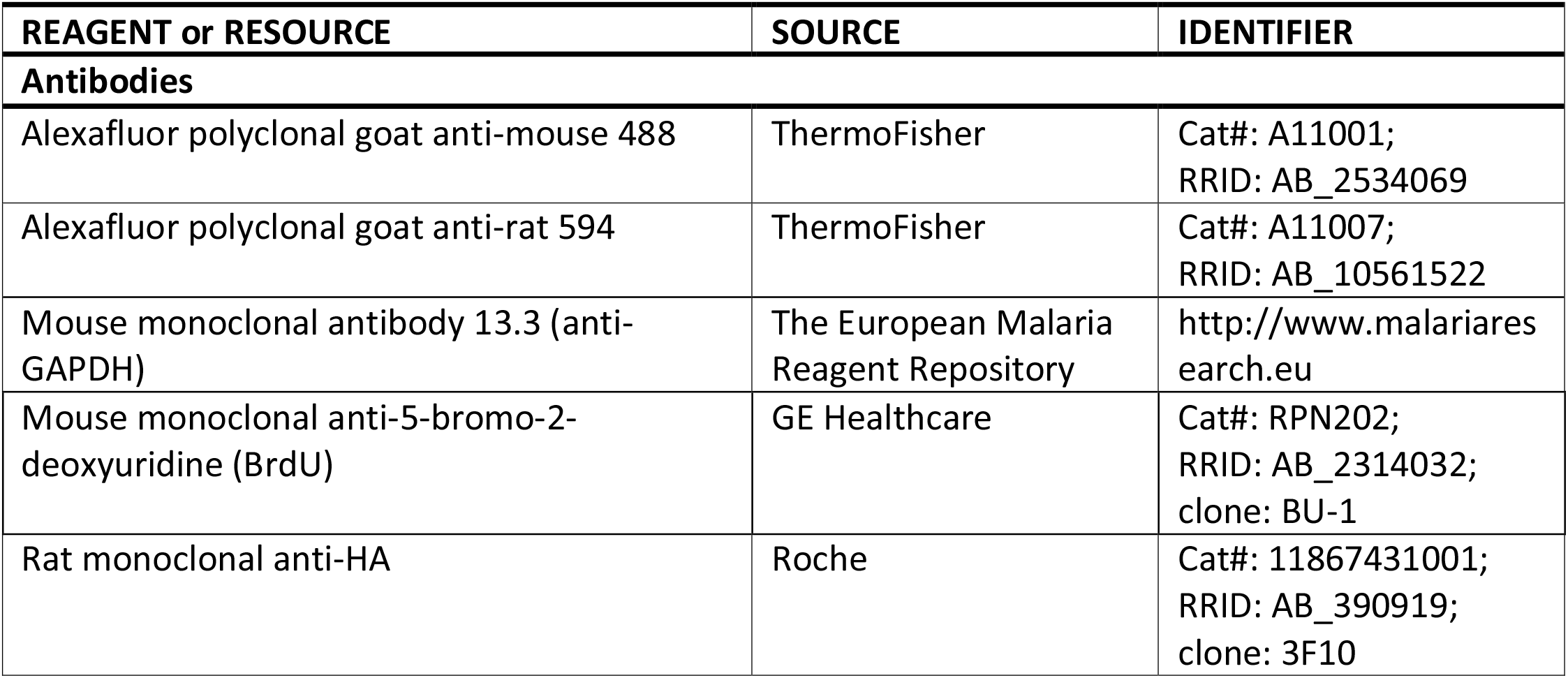
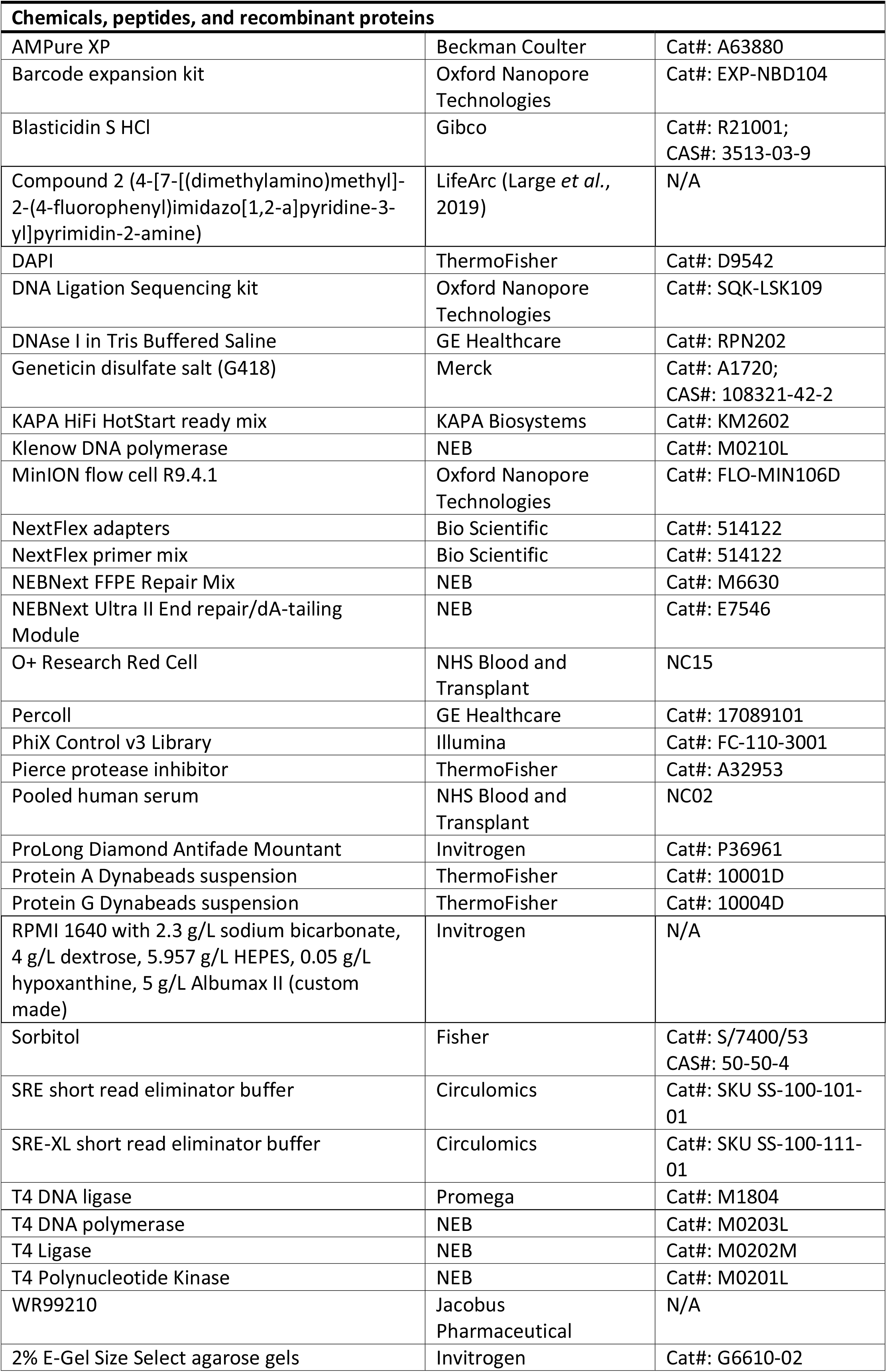
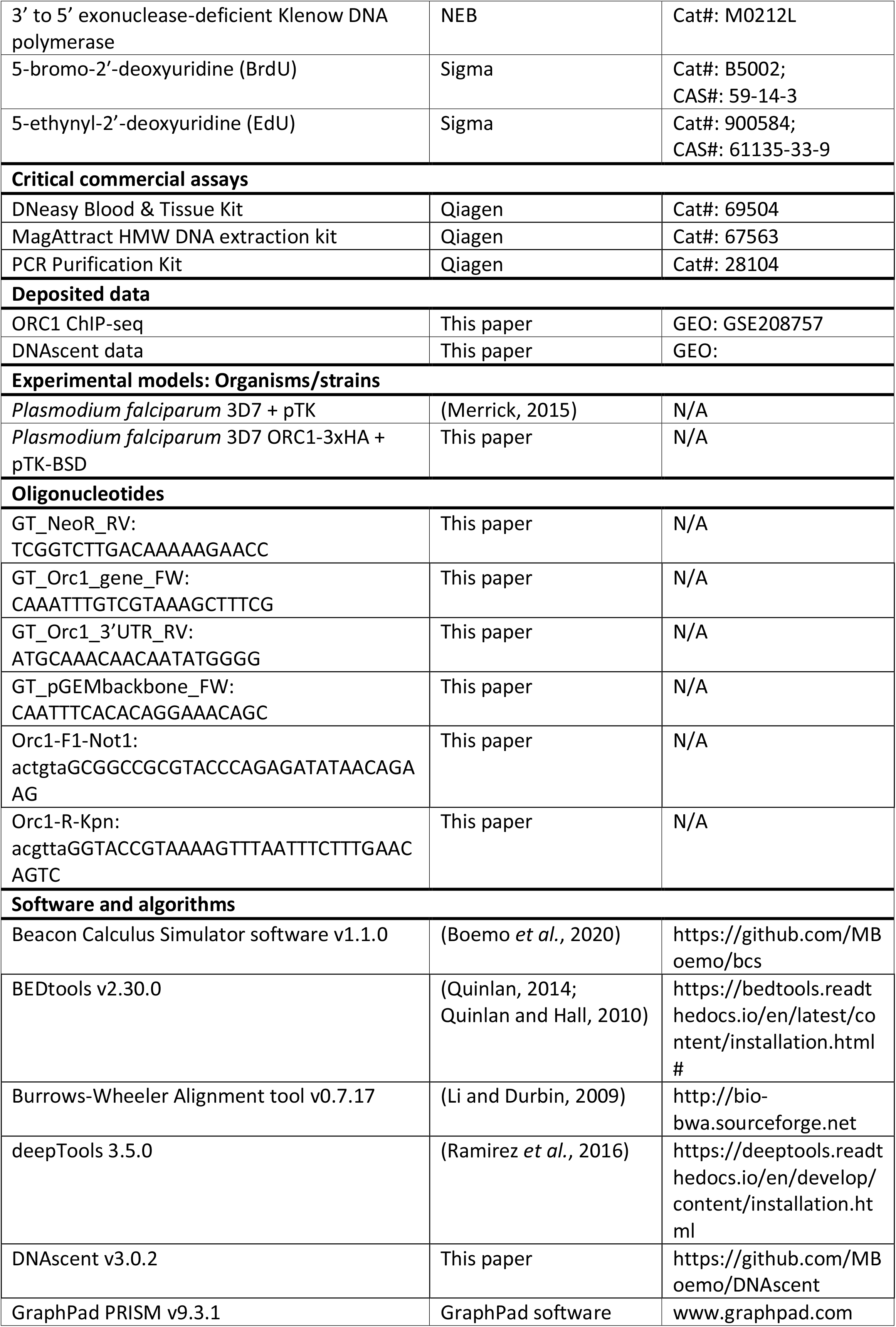
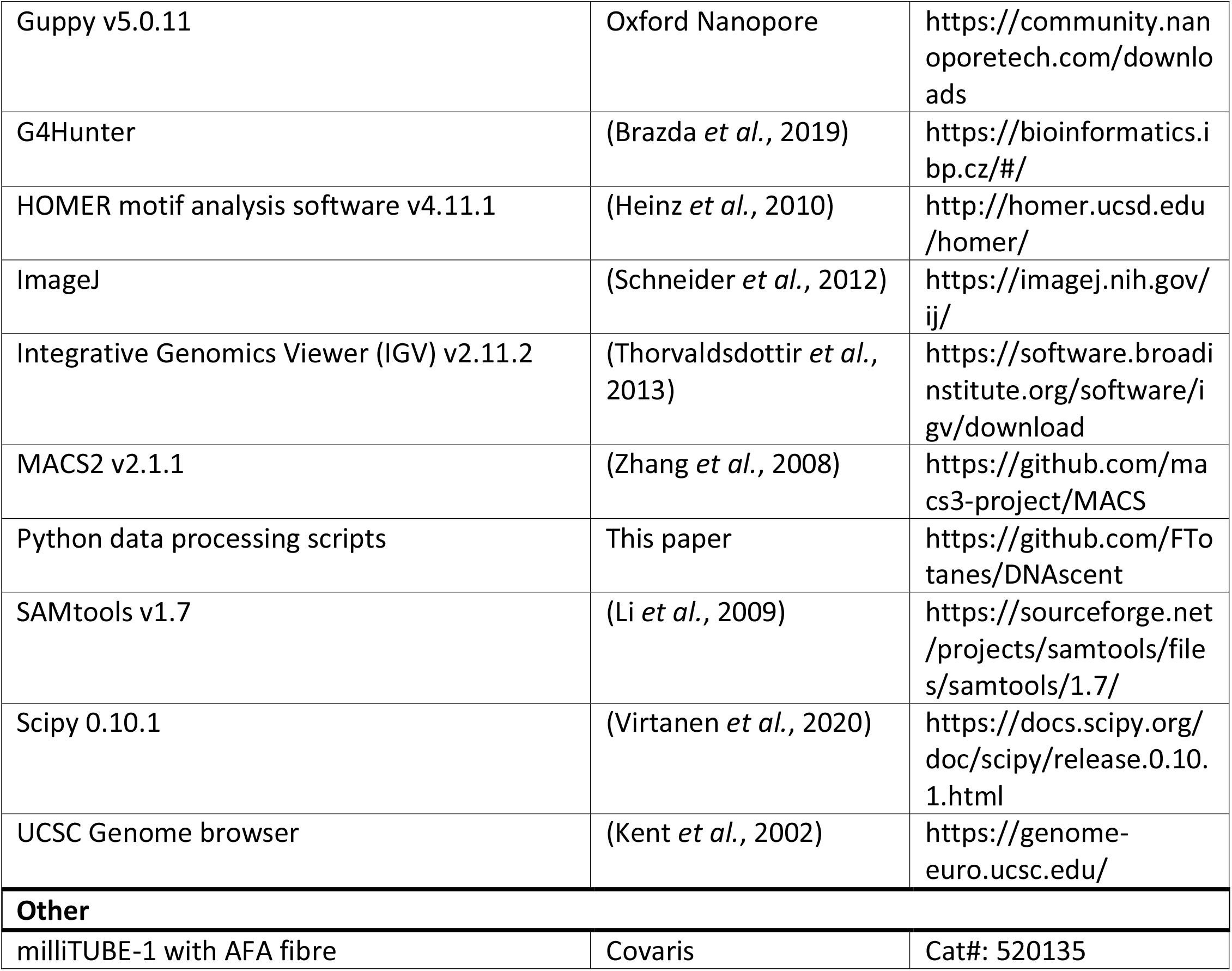

### RESOURCE AVAILABILITY

#### Lead contact

Further information and requests for resources and reagents should be directed to and will be fulfilled by the Lead Contact, Catherine Merrick.

#### Materials availability

Plasmid (pTK-BSD) and transgenic parasite line (*P. falciparum* 3D7 ORC1-3xHA + pTK-BSD) used in this study may be available upon request through the Lead Contact.

#### Data and code availability

ChIP-seq data have been deposited at GEO. DNAscent v3.0.2 is available at https://github.com/MBoemo/DNAscent and relevant Python scripts are available at https://github.com/FTotanes/DNAscent. Accession numbers are listed in the key resources table. Any additional information required to reanalyse the data reported in this paper is available from the Lead Contact upon request.

## EXPERIMENTAL MODEL

### Malaria parasites

#### Parasite culture and synchronisation

Continuous *P. falciparum* 3D7 cultures were grown in custom made RPMI 1640 (with 2.3 g/L sodium bicarbonate, 4 g/L dextrose, 5.957 g/L HEPES, 0.05 g/L hypoxanthine, 5 g/L Albumax II, Invitrogen) and 5% pooled human serum (i.e. complete media) in 4% haematocrit O+ human red blood cells in 3% oxygen, 5% CO_2_ and 92% nitrogen gas mixture at 37°C. Parasites were initially synchronised with 5% sorbitol (w/v in water) and were allowed to grow to late schizont stage. Schizonts were harvested using a 65% Percoll (GE Healthcare) gradient (v/v in PBS) and incubated for 2 hours in complete media with 1.5 μM Compound 2 (LifeArc) (Large *et al*., 2019). Compound 2 was washed off with RPMI and mature schizonts were allowed to reinvade in 25% haematocrit red blood cells in 5 ml of complete media in the abovementioned gas mixture at 37°C for 1 hour at 220 rpm. The remaining schizonts were removed using 5% sorbitol to produce a tightly synchronised culture (0 to 1 hours post invasion, referred to as 0 hpi).

#### Generation of genetically modified parasites

*P. falciparum* 3D7 *orc1* (PF3D7_1203000) was C-terminally tagged with 3xHA using the selection linked integration (SLI) technique (Birnbaum *et al*., 2017). Approximately 500 bp from the 3’ end of the *orc1* gene (excluding the stop codon) was amplified and cloned between the NotI and KpnI sites of a pSLI 3’ HA-Neomycin resistance tagging vector with a human DHFR selection marker (Supplementary Figure 2a). The plasmid (100 μg) was electroporated into a predominantly ring-stage culture and was selected initially with 5 μM WR99210 (Jacobus Pharmaceutical) followed by 400 μg/mL of G418 (Merck). Successful transfection and gene tagging was confirmed by PCR and western blot (Supplementary Figure 3a and b, Supplementary Table 7). A stable line with mostly ring-stage parasites was transfected with 100 μg of blasticidin-selectable plasmid with a thymidine kinase expression cassette (pTK-BSD) to allow episomal thymidine kinase expression and incorporation of nucleotide analogues during DNA replication (Supplementary Figure 2b). Successful transfection was selected with 2.5 μg/mL of blasticidin (Gibco) and was confirmed by ELISA and immunofluorescence for the presence of incorporated BrdU in parasite DNA (Merrick, 2015). The resulting modified parasite line, i.e. *P. falciparum* 3D7 ORC1-3xHA + pTK-BSD, was utilised for subsequent ChIP and immunofluorescence experiments. Crude parasite protein lysates from tightly synchronised parasites were fractionated into cytosolic, nuclear soluble and nuclear insoluble fractions as previously described in Voss *et al*. 2002 (Voss *et al*., 2002). Briefly, saponin-lysed parasites were incubated for 5 minutes in ice-cold lysis buffer containing 20 mM HEPES, pH 7.8, 10 mM KCl, 1 mM EDTA, 1 mM DTT, 1 mM PMSF, 0.65% Nonidet P-40. Cytoplasmic protein fraction (supernatant) was obtained by centrifugation at 2500g for 5 minutes. The pellet was then incubated in extraction buffer containing 20 mM HEPES, pH 7.8, 800 mM KCl, 1 mM EDTA, 1 mM DTT, 1 mM PMSF, 1x Pierce protease inhibitor (ThermoFisher) shaking at 2000 rpm at 4°C for 30 minutes. Centrifugation was done at 13,000g for 30 minutes to obtain the soluble and insoluble nuclear proteins (supernatant and pellet, respectively). Western blot of the fractionated samples was probed using rat anti-HA (Roche) antibodies diluted 1:1000 in block (1% BSA with 0.1% Tween 20). As control, the western blot membrane was also probed using mouse monoclonal anti-*P. falciparum* GAPDH antibody was obtained from The European Malaria Reagent Repository.

## METHOD DETAILS

### Immunofluorescence

Tightly synchronised *P. falciparum* 3D7 ORC1-3xHA + pTK-BSD culture was incubated in 100 μM BrdU (Sigma) 30 minutes prior to collection of samples for immunofluorescence staining. Parasite smears were done every 4 hours from 18 hpi to 46 hpi. Thick smears were fixed with 4% formaldehyde in PBS for 10 minutes followed by permeabilization in 0.2% Triton-X100 for 15 minutes. Slides were blocked in 1% BSA with 0.1% Tween-20 for 1 hour. Primary antibody labelling was done for 1 hour using mouse anti-BrdU (GE Healthcare) and rat anti-HA (Roche) antibodies diluted 1:500 in 1 U/mL DNAse I in Tris Buffered Saline containing 1% BSA (GE Healthcare). Three 5-minute washes with block (1% BSA with 0.1% Tween-20) were done prior to incubation in secondary antibodies (ThermoFisher Alexafluor goat anti-mouse 488 and Alexafluor goat anti-rat 594) in block at 1:1000 dilution for 1 hour. Three 5-minute final washes were done with block, with the second wash replaced with DAPI (ThermoFisher) at 2 μg/mL in PBS. All incubation steps were done at room temperature. Slides were cured using ProLong Diamond Antifade Mountant (Invitrogen) and were stored at 4°C prior to visualisation.

Images were acquired using a Nikon Microphot SA microscope with a Qimaging Retiga R6 camera at 1000x magnification. ImageJ (Schneider *et al*., 2012) was used to identify and create the corresponding regions of interest (ROIs) on nuclear signal in the DAPI channel images. These ROIs where then used to measure the area and integrated signal density in all images taken using the different fluorescent channels. The resulting data were analysed and plotted using GraphPad Prism v9.3.1. Representative images were pseudo-coloured and merged using ImageJ.

### Chromatin immunoprecipitation

#### Chromatin preparation

Cultures of 2.0 × 10^9^ and 2.4 × 10^9^ tightly synchronized parasites at 24 and 30 hpi, respectively, were used for ChIP. Chromatin was crosslinked with 1% formaldehyde in culture media for 10 minutes at 37°C, then quenched with glycine at a final concentration of 0.125 M on ice. Samples from here on were maintained on ice. Parasites were extracted by lysis with 0.05% saponin in PBS. Nuclei were extracted by gentle homogenisation in cell lysis buffer (10 mM Tris pH 8.0, 3 mM MgCl_2_, 0.2% NP-40, 1x Pierce protease inhibitor (ThermoFisher)) and centrifugation at 2000 rpm for 10 minutes in 0.25 M sucrose cushion in cell lysis buffer. Harvested nuclei were snap-frozen in 20% glycerol in cell lysis buffer. Samples were resuspended in shearing buffer (Tris-HCl 10 mM, EDTA 1 mM, 0.1% SDS, pH 7.6) to a final volume of 1 mL and sonicated in a milliTUBE-1 with AFA fibre (Covaris) using a Covaris M220 sonicator with the following settings:

Duty factor: 5%

Temperature: min: 5.0°C; setpoint: 7.0°C; max: 9.0°C

Peak power: 75.0 W

Cycle/burst: 200

Processing time: 2700 sec (45 min)

#### Chromatin immunoprecipitation

Each ChIP reaction was set up with 500 ng sonicated chromatin in incubation buffer (0.15% SDS, 1% Triton-X100, 150 mM NaCl, 1 mM EDTA, 0.5 mM EGTA, 1x protease inhibitor (Sigma-Aldrich), 20 mM HEPES, pH 7.4) with 400 ng of anti-HA (Roche), together with 10 μL protA and 10 μL protG Dynabeads suspension (ThermoFisher). For each sample, eight ChIP reactions were prepared and incubated overnight rotating at 4°C. Beads were washed twice with wash buffer 1 (0.1% SDS, 0.1% DOC, 1% Triton-X100, 150 mM NaCl, 1 mM EDTA, 0.5 mM EGTA, 20 mM HEPES, pH 7.4), once with wash buffer 2 (0.1% SDS, 0.1% DOC, 1% Triton-X100, 500 mM NaCl, 1 mM EDTA, 0.5 mM EGTA, 20 mM HEPES, pH 7.4), once with wash buffer 3 (250 mM LiCl, 0.5% DOC, 0.5% NP-40, 1 mM EDTA, 0.5 mM EGTA, 20 mM HEPES, pH 7.4) and twice with wash buffer 4 (1 mM EDTA, 0.5 mM EGTA, 20 mM HEPES, pH 7.4). Each wash step was performed for 5 minutes rotating at 4°C. Immunoprecipitated chromatin was eluted in elution buffer (1% SDS, 0.1M NaHCO_3_) at room temperature for 20 min. The eluted chromatin samples and the corresponding input samples (sonicated input chromatin containing 500 ng DNA) were de-crosslinked in 1% SDS / 0.1M NaHCO_3_ / 1M NaCl at 45°C for overnight while shaking, followed by column purification (PCR Purification Kit, Qiagen) and elution in 20 μL EB buffer.

The obtained ChIP-ed DNA fragments were used to generate Illumina sequencing libraries according to Filarsky *et al*. (Filarsky *et al*., 2018). In brief, 5 ng of α-HA ChIP or input DNA were end-repaired with T4 DNA polymerase (NEB), Klenow DNA polymerase (NEB), and T4 Polynucleotide Kinase (NEB). The 3’ends of end-repaired DNA were extended with an A-overhang with 3’ to 5’ exonuclease-deficient Klenow DNA polymerase (NEB). The resulting fragments were ligated to Nextflex adapters (Bio Scientific) with the use of T4 DNA ligase (Promega). The libraries were amplified using an AT-rich optimized KAPA protocol using KAPA HiFi HotStart ready mix (KAPA Biosystems), NextFlex primer mix (Bio Scientific) with the following PCR program: 98°C for 2 min; four cycles of 98°C for 20 sec, 62°C for 3 min; 62°C for 5 min. The fragments originating from mono-nucleosomes + 125 bp NextFlex adapter were selected using 2% E-Gel Size Select agarose gels (Invitrogen) and amplified by PCR for 8 cycles using the above conditions. Libraries were purified and adapter dimers removed with Agencourt AMPure XP beads purification using a 1:1 library:beads ratio (Beckman Coulter). ChIP-seq libraries were sequenced on the Illumina NextSeq 500 system with a 20% phiX spike-in (Illumina) to generate 75 bp single-end reads (NextSeq 500/550 High Output v2 kit).

Sequencing reads were mapped against the PlasmoDB v26 *P. falciparum* genome utilising the Burrows-Wheeler Alignment tool (v0.7.17) (Li and Durbin, 2009). Mapped reads were filtered to mapping quality ≥ 30 (SAMtools v1.7) (Li *et al*., 2009) and only uniquely mapped reads were used for further analysis.

### Datasets for EdU and BrdU model training

To produce training data sets for the DNAscent algorithm, tightly synchronised thymidine kinase-expressing *P. falciparum* 3D7 culture (Merrick, 2015) was incubated with either 20 μM EdU (ThermoFisher) or 200 μM BrdU (Sigma) for 1 hour. Parasite DNA was then harvested using a DNeasy Blood and Tissue DNA extraction kit (Qiagen). DNA barcoding was done on 1 μg of high molecular weight genomic DNA using Oxford nanopore ligation sequencing kit (SQK-LSK109) and barcode expansion kit (EXP-NBD104). Sequencing was done using a nanopore MinION device with R9.4.1 flowcells for 72 hours or until the pores in the flowcell have been fully depleted. To produce data sets for the identification of active replication forks and origins, tightly synchronised parasites were incubated in 20 μM EdU for 7.5 minutes, then 200 μM BrdU for another 7.5 minutes, followed by 2 mM thymidine for 1 hour and 45 minutes prior to parasite harvest. Nascent DNA labelling was done at 30 and 36 hpi. High molecular weight parasite genomic DNA was extracted using a Qiagen MagAttract kit and stored at 4°C. Long DNA fragments were enriched using SRE/SRE-XL short read eliminator buffer (Circulomics) prior to library preparation and nanopore sequencing.

## QUANTIFICATION AND STATISTICAL ANALYSIS

### DNAscent software development and model training

Given the stochasticity of DNA replication in erythrocytic schizogony, we expected only a minority of sequenced reads in the training set to be analogue-substituted. While DNAscent v2.0.2 (Boemo, 2021) was trained to identify BrdU, we found that the current shifts caused by BrdU and EdU were sufficiently similar that DNAscent v2.0.2 often called EdU as BrdU in EdU-substituted DNA (Supplementary Figure 4). DNAscent v2.0.2 was therefore used to identify and pull out analogue-substituted reads that were suitable for training. Reads where DNAscent v2.0.2 called BrdU with probability greater than 0.5 for at least 20% of thymidine positions were considered to be analogue-substituted, resulting in 6,917 BrdU-substituted reads and 3,162 EdU-substituted reads. Training reads were split into 4 kb segments and augmented with thymidine reads as in (Boemo, 2021). The neural network architecture used was identical to that of (Boemo, 2021), with the exception that the dimension of the time-distributed softmax layer in was increased from two to three to reflect the three classification categories (thymidine, BrdU, or EdU). The model was trained for 21 epochs on 15,000 EdU-augmented segments, 15,000 BrdU-augmented segments, and 10,000 thymidine-only segments. ROC curves were computed in *S. cerevisiae* reads from (Muller *et al*., 2019) to show that the new model, despite calling two thymidine analogues instead of one, had similar BrdU-calling accuracy and lower false positive rate compared to the BrdU-only model used in DNAscent v2.0.2 (Supplementary Figure 5). For the new model, ROC curves computed on *P. falciparum* test reads showed BrdU- and EdU-calling performance were similar to each other, the frequency with which the model would confuse BrdU for EdU (and vice versa) was low, and that the model had a very low false positive rate of approximate 1-in-700 thymidines (Supplementary Figure 6).To make replication origin, fork, and termination calls from patterns of analogue calls, the rate analogue incorporation in analogue-substituted regions was measured by K-means clustering (K=2) the rate of analogue substitution in consecutive 2 kb segments. The rate of analogue incorporation in analogue-substituted regions was taken to be the greater of the two centroid means. Reads were then segmented with a DBSCAN algorithm (epsilon=1000 bp; minimum density=analogue centroid mean – 1 standard deviation). Different analogue segments closer than 5 kb apart were matched into forks and origins were called as the genomic regions between matched left- and rightward-moving forks.

### Data analysis

ChIP and DNAscent origin log2ratios were calculated using deepTools 3.5.0 bamCompare by normalising reads against input/background reads, with a pseudocount of +1 and a bin size of 50 bp (Ramirez *et al*., 2016). Normalisation was done separately for the two ChIP replicates. Nanopore read normalisation was done on a per barcode basis and log2ratios were summed by timepoint. MACS2 v2.1.1 was used for ChIP peak calling with p-value cut-off of 1e-3 (Zhang *et al*., 2008). BEDtools v2.30.0 (Quinlan, 2014) was used for file format conversions, comparison of multiple data sets, and identifying genes overlapping with areas of interest in the genome (bedtools intersect). GC-bias (computeGCBias), Spearman R correlation analyses between multiple bigwig files (multiBigwigSummary, bin size of 100 bp), and matrix plots (computeMatrix) were done using deepTools 3.5.0. Fork speed calculation were done on all forks identified by DNAscent that are not located at the nanopore read ends, or joint at the replication origin or termination site. Genome-wide mean fork speed z-score per 100 bp bin was calculated as a metric to determine the deviation of the mean fork speed at any given region in the genome from the overall mean fork speed. Fork speed was calculated by dividing the fork length (in kb) by 15 minutes (the duration of the pulse) giving the fork speed in kb/min. Fork speed z-score was then calculated using the formula:

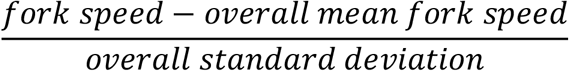

Fork speed z-score per 100 bp was determined using bedtools closest to identify forks that overlap any given 100 bp window in the genome, and bedtools merge to calculate the mean per window.

*De novo* motif search was done on ORC1 ChIP summits ± 50 bp using HOMER motif analysis software v4.11.1 (Heinz *et al*., 2010). Search parameters included motif lengths of 8, 10, 12, 15, 20, 25 and 30 bp and sequence content G/C normalisation. The calculation of hypergeometric p-values was done in reference to motif occurrence in preparsed background sequences. Similar motif search parameters were used on DNAscent origin centres ± 50 bp. Motifs with p-values ≤ 1e-12 were considered significant. Motif weighted G/C content was calculated as the sum of the individual probability of a G or C in the motif sequence divided by the sum of the probabilities for all nucleotides. G4Hunter (Brazda *et al*., 2019) was used to identify G-quadruplex motifs in ORC1 ChIP peaks with the following parameters: window size of 25 and threshold of 1.7. Enrichment in ChIP peak sequences was calculated as the total number of identified G4 per bp divided by the total number of G4 (+1, to avoid a zero denominator) per bp in the entire genome.

Data on predicted AP2 transcription binding sites (Campbell *et al*., 2010) and genome-wide HP1 log2ratio (Fraschka *et al*., 2018) were downloaded from PlasmoDB (Aurrecoechea *et al*., 2009). Other data sets utilised in the analyses were RNA-seq/ATAC-seq (Toenhake *et al*., 2018) and histone placements (Bartfai *et al*., 2010). Gene ontology (GO) terms were obtained from PlasmoDB (Aurrecoechea *et al*., 2009) and were collated and summarised using Python. Bigwig, bedgraph and bed files were visualised using Integrative Genomics Viewer (IGV) version 2.11.2 (Thorvaldsdottir *et al*., 2013) or UCSC Genome browser (Kent *et al*., 2002). D’Agostino and Pearson’s test for normality, Mann Whitney U test for significant difference, and Spearman R test for correlation between smaller data sets were done using Scipy 0.10.1 (Virtanen *et al*., 2020).

To call replication forks and origins on single molecules, Oxford nanopore sequencing reads were basecalled and demultiplexed with Guppy (v5.0.11) and aligned to the *P. falciparum* 3D7 ASM276v2 assembly (https://www.ncbi.nlm.nih.gov/assembly/GCF_000002765.4/) with minimap2 (v2.17-r941). Only sequences with an alignment length ≥ 10 kb and mapping quality ≥ 20 were analysed. The probability of BrdU and EdU at thymidine positions along each read was assigned by DNAscent detect, and these probabilities were parsed into replication fork and origin calls by DNAscent forkSense. Bedgraphs of base analogue calls on single molecules were generated using the dnascent2bedgraph utility. DNAscent detect, DNAscent forkSense, and dnascent2bedgraph are all part of DNAscent v3.0.2 available under GPL-3.0 at https://github.com/MBoemo/DNAscent.

## SUPPLEMENTAL ITEM TITLES

**Supplementary Figure 1:**
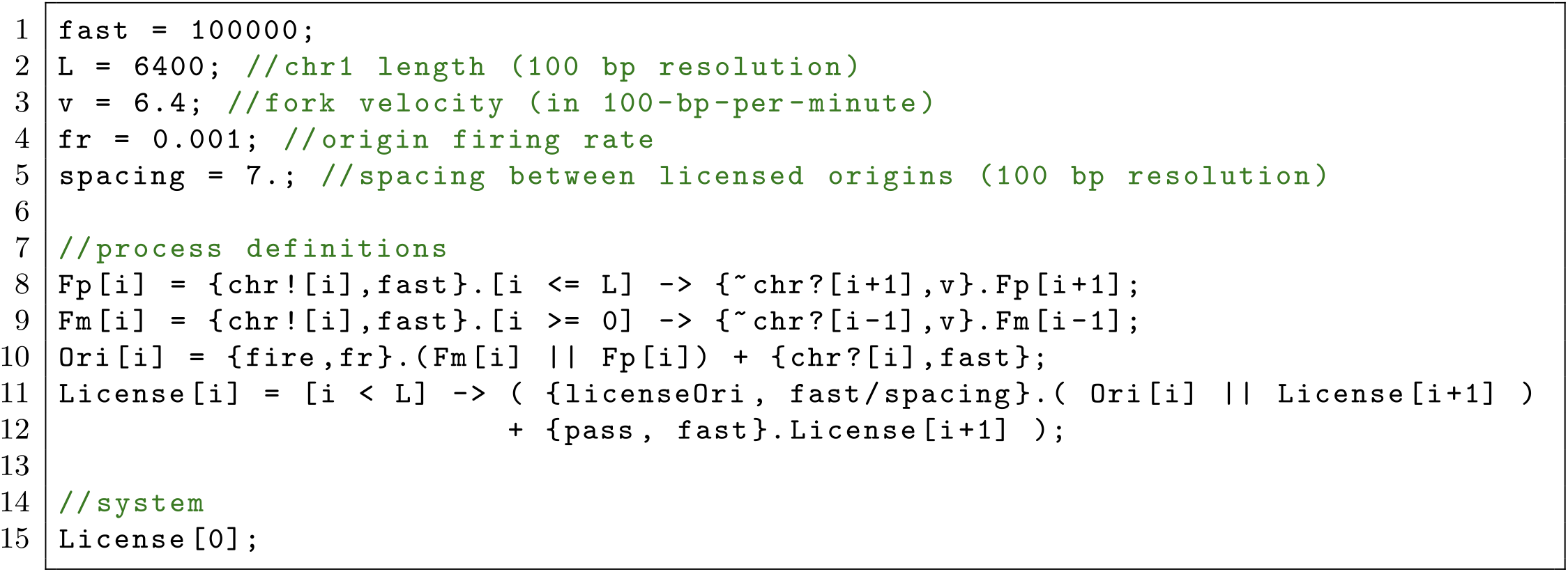
Source code of the Beacon Calculus model of the DNA replication of *P. falciparum*. Source code of the Beacon Calculus model of the DNA replication of *P. falciparum* chromosome 1 with 100-bp resolution. The model includes parameters for the length of *P. falciparum* chromosome 1 (Line 2), a uniform rate of fork movement (Line 3), a uniform firing rate for all origins (Line 4), and an average spacing between licensed origins (Line 5). The fork processes (Lines 8-9) and the origin process (Line 10) behave as in (Boemo *et al*., 2020). An origin licensing process (Lines 11-12) distributes licensed origins across the chromosome with an average spacing of 700 bp.

**Supplementary Figure 2:**
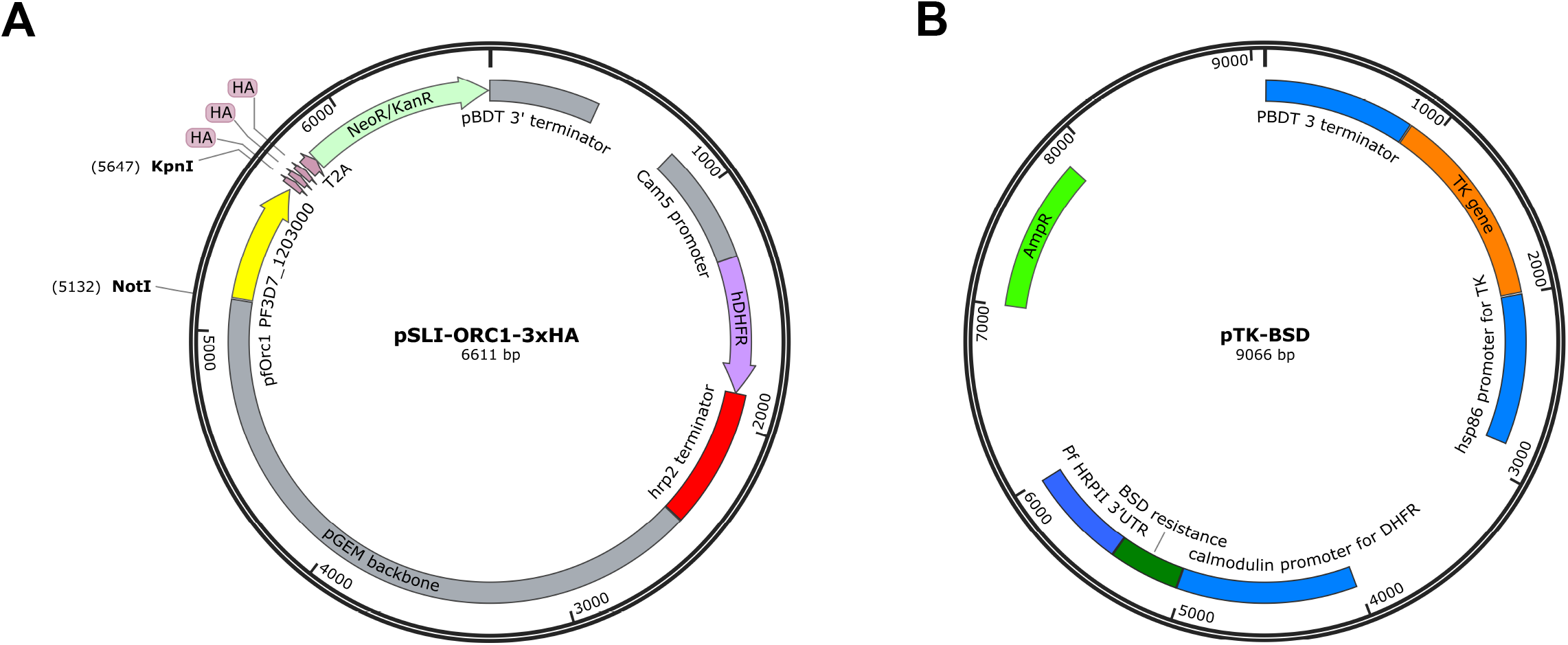
Plasmids used in the generation of genetically modified parasites. A) Selection-linked integration plasmid used to C-terminally tag *P. falciparum* 3D7 with 3xHA. B) Transfected plasmid used to allow parasites to episomally express thymidine kinase.

**Supplementary Figure 3:**
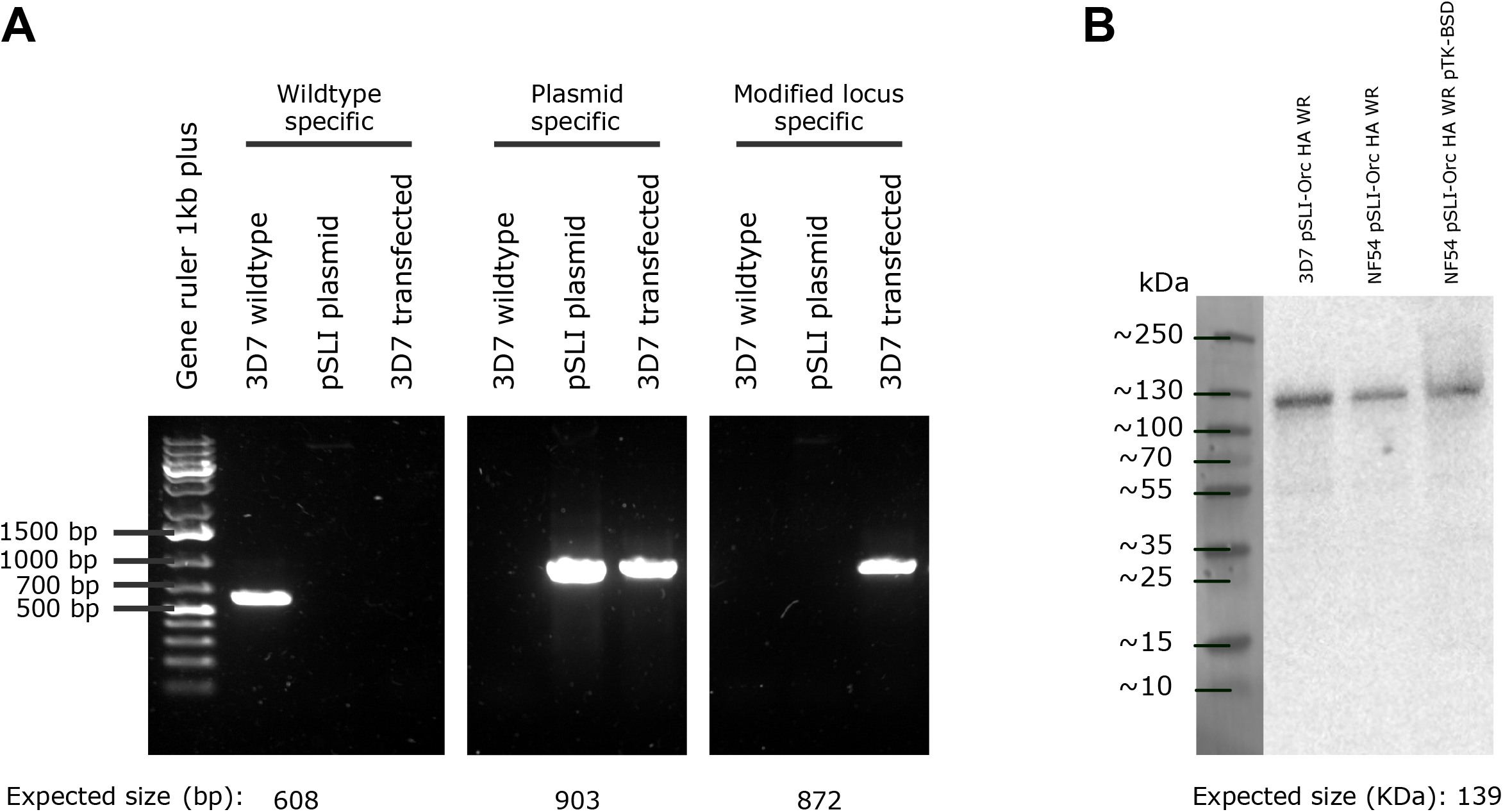
Confirmation of transfection and gene tagging. A) PCR confirmation of successful integration of 3xHA and neomycin resistance gene into *orc1* (PF3D7_1203000). Primers used are listed in Supplementary Table 7 and Key Resources Table. B) Western blot confirmation of HA-tagging of ORC1.

**Supplementary Figure 4:**
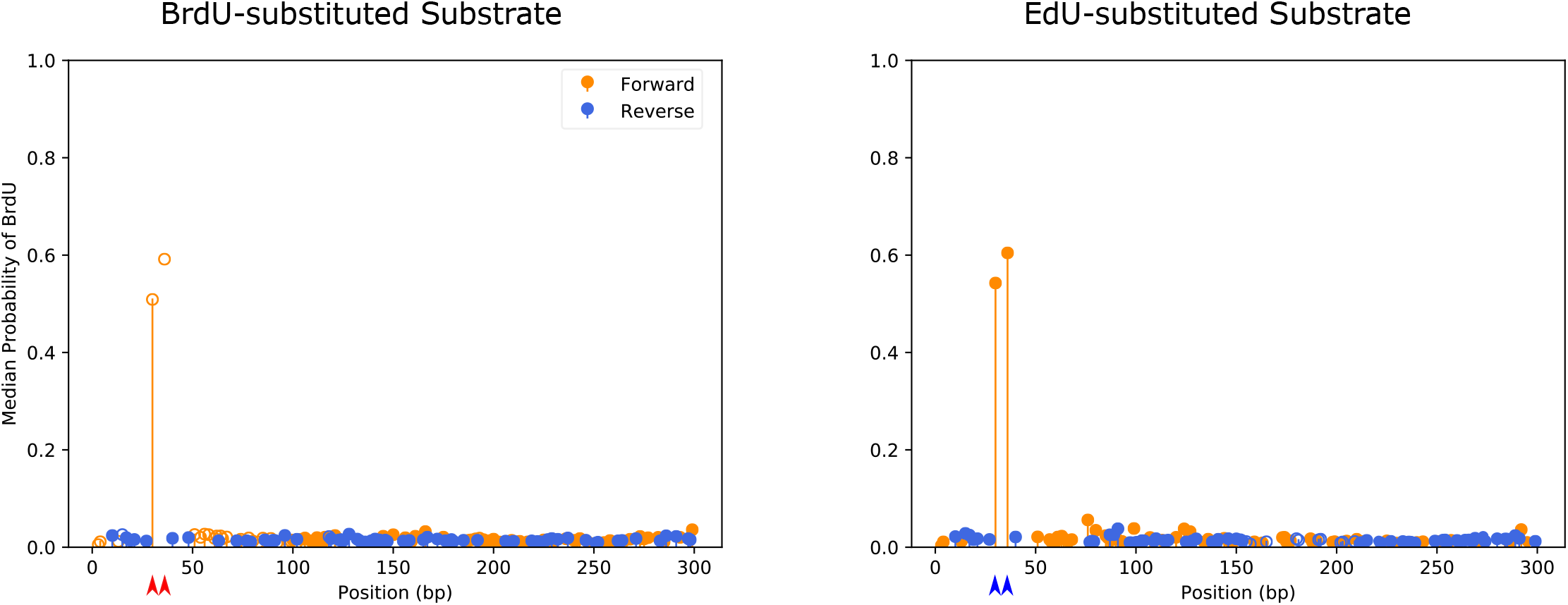
Median probabilities of BrdU and EdU called using DNAscent v2.0.2. Median probabilities of BrdU called using DNAscent v2.0.2 on the forward (blue) and reverse (orange) strands of primer extension reads from (Muller *et al*., 2019) where either BrdU or EdU were incorporated into two known positions (30 and 36 bp; red arrows for BrdU; blue arrows for EdU) on the forward strand. Reads analysed had a mapping length greater than or equal to 100 bp and mapping quality greater than or equal to 20 were used, resulting in N=453 reads for the BrdU-substituted substrate (left) and N=594 reads for the EdU-substituted substrate (right).

**Supplementary Figure 5:**
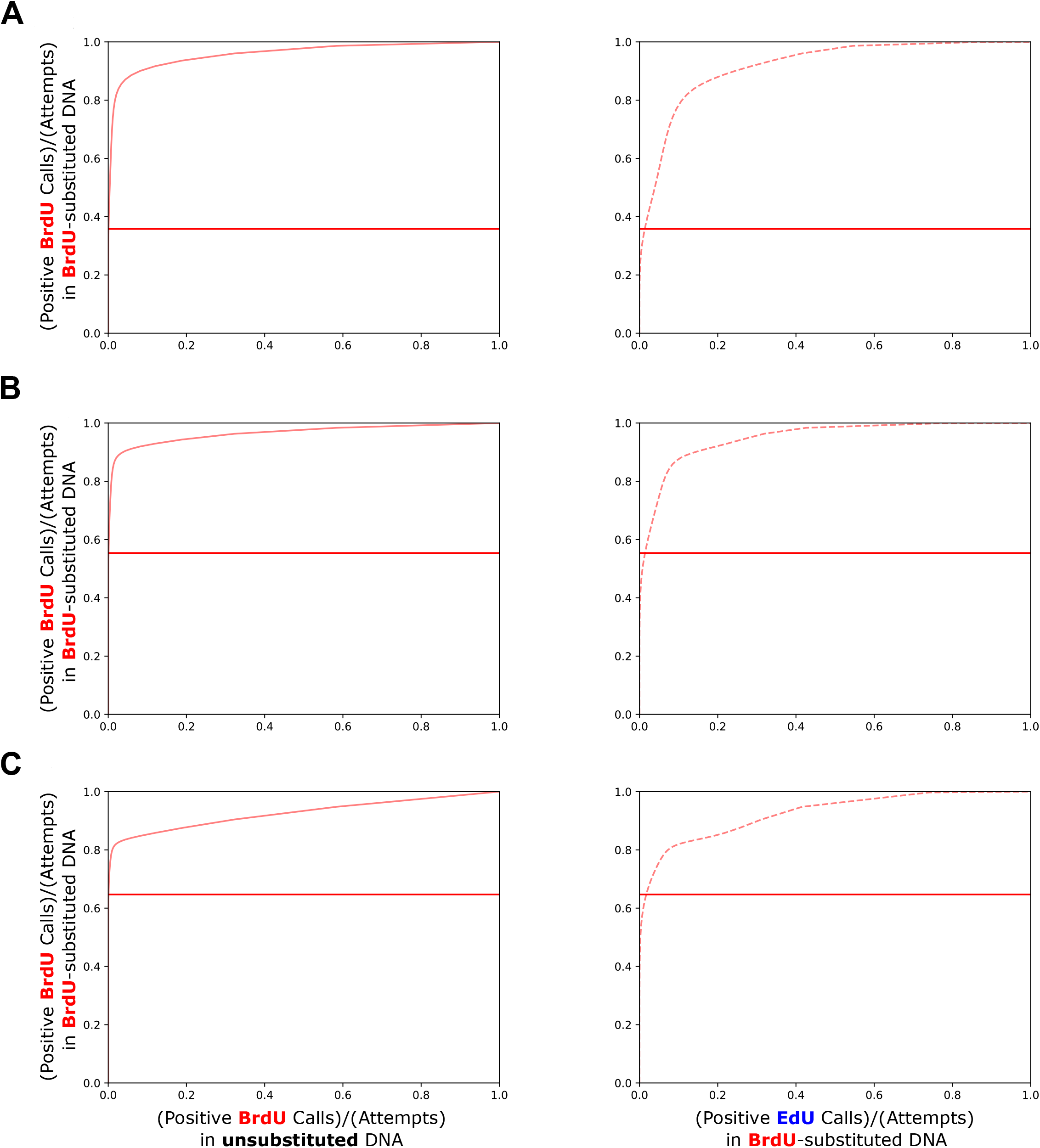
Receiver operator characteristic curves for *S. cerevisiae* reads. Receiver operator characteristic (ROC) curves for *S. cerevisiae* reads from (Muller *et al*., 2019) where BrdU substitution was measured by mass spectrometry to find (A) 26%, (B) 49%, and (C) 79% substitution. Curves show the number of positive analogue calls, defined as thymidine positions where DNAscent v3.0.2 assigned a probability of that analogue (BrdU or EdU) greater than a probability threshold, divided by number of thymidine positions. Points along the ROC curves indicate different probability thresholds above which an analogue call is considered positive. Horizontal lines intersect the curves at probability threshold of 0.5. Each curve was computed using N=1000 reads with a mapping quality of at least 20 and a mapping length of at least 1 kb to the *S. cerevisiae* sacCer3 assembly.

**Supplementary Figure 6:**
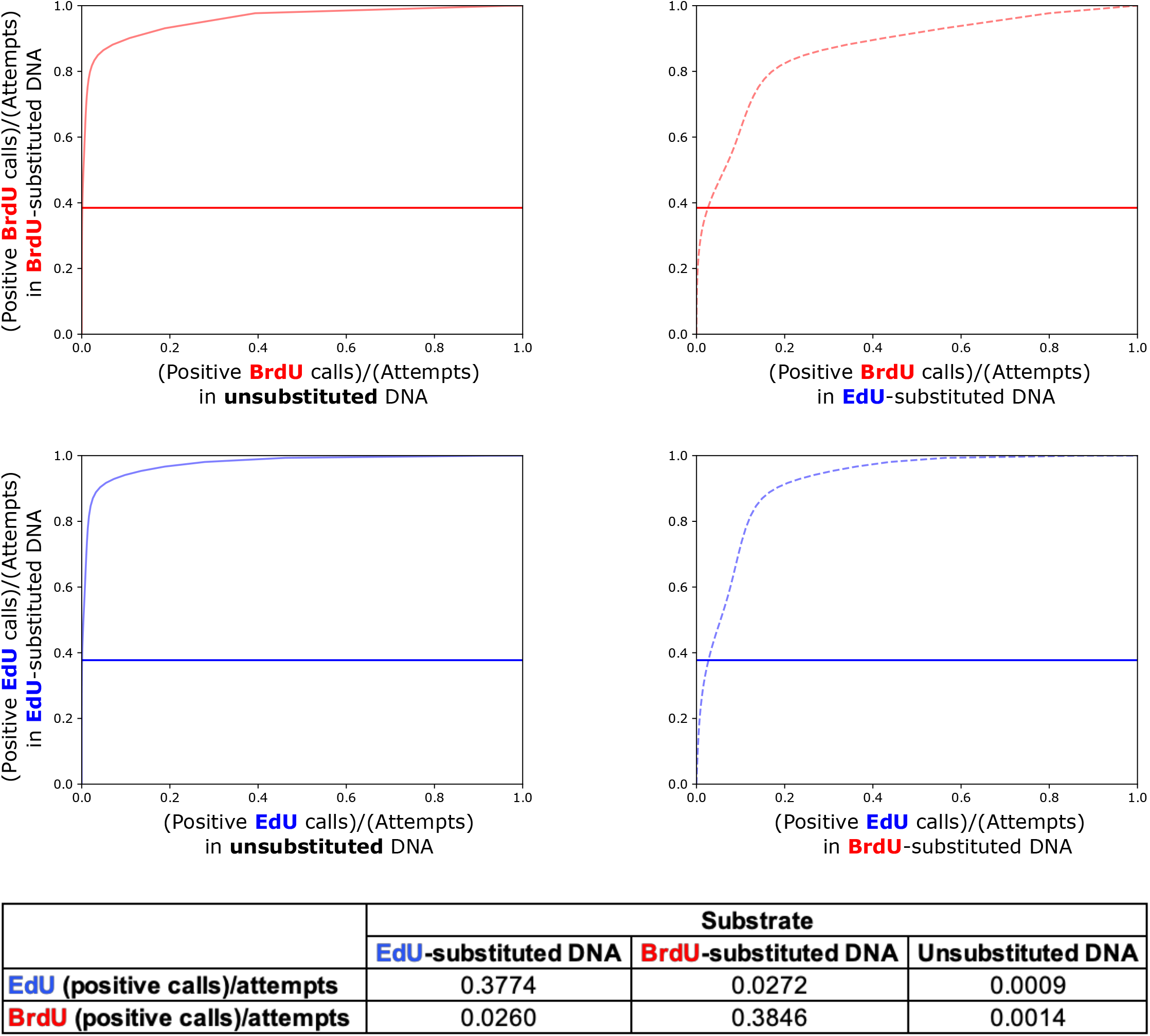
Receiver operator characteristic curves for *P. falciparum* reads. Receiver operator characteristic (ROC) curves for *P. falciparum* reads, where the probability of BrdU and EdU at each thymidine position was measured by DNAscent v3.0.2. The analogue-substituted reads for this benchmark were taken from a biological repeat of the sequencing run used to generate training data, and hence these reads are independent of the training dataset. As in the training dataset, EdU- and BrdU-treated *P. falciparum* reads were given distinct barcodes, sequenced on the Oxford nanopore platform, and demultiplexed using Guppy. Reads were considered analogue-substituted if DNAscent v2.0.2 called made positive BrdU calls (probability of BrdU > 0.5) in at least 20% of thymidine positions across the read. Similar to Supplementary Figure 5, each curve was computed using 1000 reads with a mapping quality of at least 20 and a mapping length of at least 1 kb to the *P. falciparum* 3D7 ASM276v2 assembly and points along the ROC curves show probability thresholds above which an analogue call is considered positive. Horizontal lines intersect the curves at probability threshold of 0.5. The table shows the values on the x- and y-axis where the horizontal line intersects the ROC curve.

**Supplementary Table 1:** ORC1 ChIP-seq inter-summit distances. Descriptive statistics of calculated distances between summits called using MACS2 on ORC1 ChIP-seq at 24 and 30 hpi per chromosome.

| Chromosome | (n) | mean | median | min | max | 10th percentile | 25th percentile | 75th percentile | 90th percentile | Chromosome length |
| --- | --- | --- | --- | --- | --- | --- | --- | --- | --- | --- |
| Pf3D7_01_v3 | 334 | 1786 | 716 | 96 | 20002 | 303 | 440 | 2177 | 4441 | 640851 |
| Pf3D7_02_v3 | 516 | 1818 | 831 | 74 | 21263 | 297 | 445 | 2520 | 4488 | 947102 |
| Pf3D7_03_v3 | 623 | 1680 | 892 | 144 | 15009 | 313 | 445 | 2187 | 4218 | 1067971 |
| Pf3D7_04_v3 | 699 | 1695 | 727 | 119 | 29594 | 296 | 433 | 2131 | 4314 | 1200490 |
| Pf3D7_05_v3 | 738 | 1787 | 906 | 86 | 18671 | 316 | 476 | 2465 | 4424 | 1343557 |
| Pf3D7_06_v3 | 781 | 1805 | 774 | 62 | 38384 | 304 | 432 | 2243 | 4659 | 1418242 |
| Pf3D7_07_v3 | 823 | 1737 | 842 | 54 | 20112 | 297 | 444 | 2275 | 4353 | 1445207 |
| Pf3D7_08_v3 | 812 | 1676 | 749 | 32 | 17052 | 302 | 430 | 2166 | 4223 | 1472805 |
| Pf3D7_09_v3 | 771 | 1964 | 960 | 77 | 26483 | 313 | 460 | 2693 | 4899 | 1541735 |
| Pf3D7_10_v3 | 927 | 1810 | 853 | 53 | 32133 | 322 | 483 | 2539 | 4385 | 1687656 |
| Pf3D7_11_v3 | 1017 | 1988 | 846 | 76 | 35943 | 318 | 462 | 2743 | 4890 | 2038340 |
| Pf3D7_12_v3 | 1256 | 1789 | 867 | 73 | 21192 | 316 | 466 | 2248 | 4591 | 2271494 |
| Pf3D7_13_v3 | 1464 | 1976 | 963 | 11 | 24713 | 310 | 471 | 2630 | 4772 | 2925236 |
| Pf3D7_14_v3 | 1645 | 2000 | 1003 | 52 | 23541 | 319 | 477 | 2784 | 4994 | 3291936 |
| Overall | 12406 | 1848 | 863 | 11 | 38384 | 310 | 457 | 2471 | 4640 | 23292622 |

30 hpi ORC1 ChIP inter-summit distances (bp)
| Chromosome | (n) | mean | median | min | max | 10th percentile | 25th percentile | 75th percentile | 90th percentile | Chromosome length |
| --- | --- | --- | --- | --- | --- | --- | --- | --- | --- | --- |
| Pf3D7_01_v3 | 271 | 2252 | 884 | 39 | 20212 | 283 | 438 | 3046 | 5362 | 640851 |
| Pf3D7_02_v3 | 393 | 2387 | 1120 | 8 | 22037 | 284 | 490 | 3424 | 5714 | 947102 |
| Pf3D7_03_v3 | 450 | 2333 | 1144 | 64 | 23039 | 343 | 562 | 3167 | 5678 | 1067971 |
| Pf3D7_04_v3 | 540 | 2200 | 832 | 58 | 30820 | 297 | 463 | 3070 | 5539 | 1200490 |
| Pf3D7_05_v3 | 530 | 2488 | 1287 | 61 | 26484 | 362 | 547 | 3337 | 5849 | 1343557 |
| Pf3D7_06_v3 | 549 | 2567 | 1112 | 111 | 39147 | 346 | 512 | 3276 | 6643 | 1418242 |
| Pf3D7_07_v3 | 585 | 2451 | 1199 | 44 | 21136 | 313 | 512 | 3433 | 6119 | 1445207 |
| Pf3D7_08_v3 | 569 | 2394 | 948 | 70 | 21905 | 315 | 483 | 3206 | 6548 | 1472805 |
| Pf3D7_09_v3 | 525 | 2885 | 1629 | 125 | 21792 | 365 | 622 | 4042 | 7061 | 1541735 |
| Pf3D7_10_v3 | 694 | 2419 | 1121 | 11 | 32303 | 316 | 506 | 3345 | 6076 | 1687656 |
| Pf3D7_11_v3 | 721 | 2789 | 1143 | 57 | 36000 | 321 | 516 | 3837 | 6764 | 2038340 |
| Pf3D7_12_v3 | 952 | 2376 | 1009 | 84 | 34858 | 314 | 478 | 3225 | 6034 | 2271494 |
| Pf3D7_13_v3 | 1033 | 2800 | 1369 | 5 | 33700 | 333 | 530 | 3882 | 6788 | 2925236 |
| Pf3D7_14_v3 | 1171 | 2809 | 1359 | 37 | 23799 | 330 | 509 | 3987 | 6998 | 3291936 |
| Overall | 8983 | 2556 | 1156 | 5 | 39147 | 323 | 509 | 3490 | 6430 | 23292622 |

**Supplementary Table 2:**
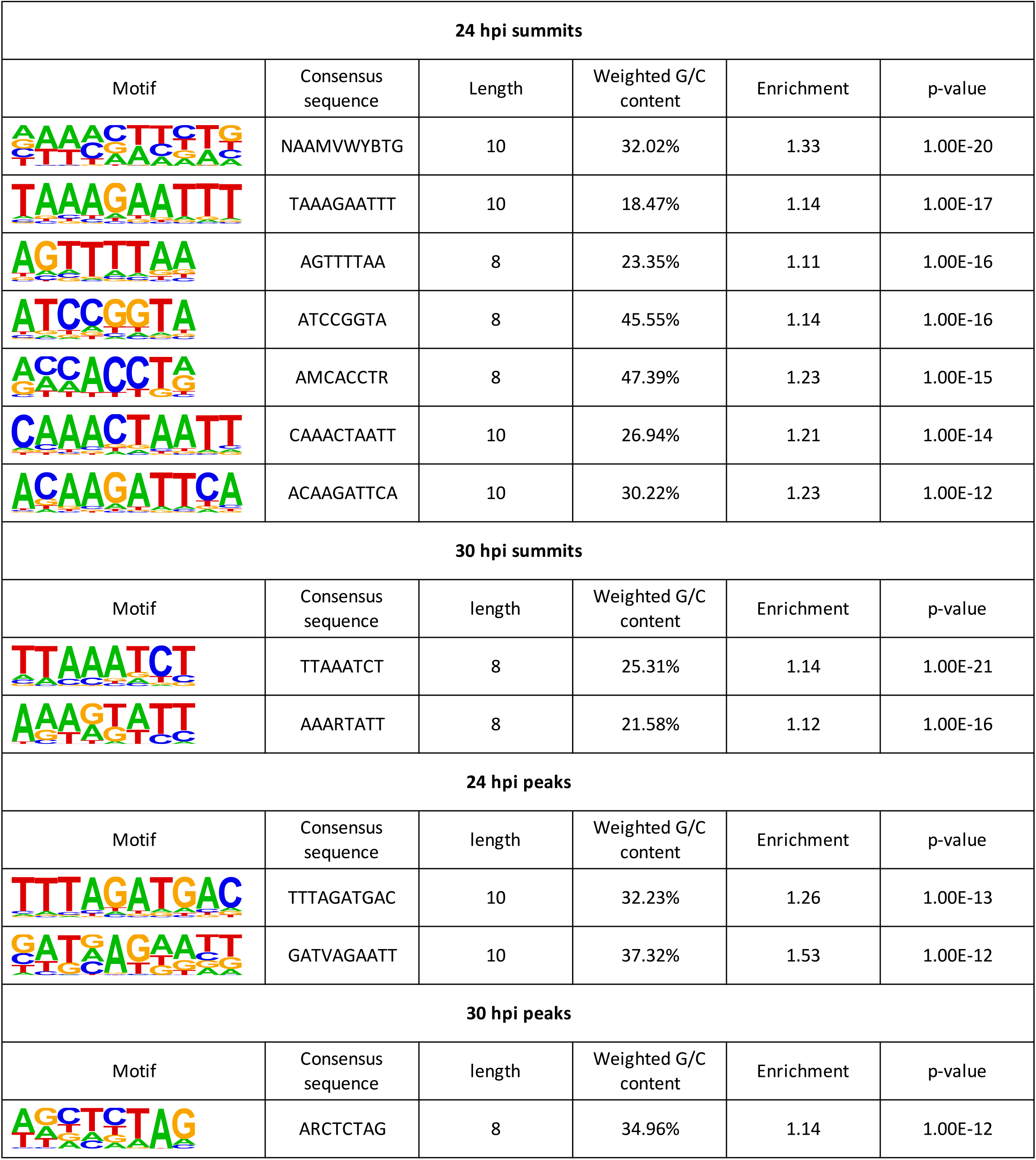
*De novo* Motifs. Significant motif structure and consensus sequences identified in ORC1 ChIP summit ± 50 bp using HOMER motif analysis software v4.11.1 with their corresponding weighted GC-content, enrichment scores and p-values. Motifs with an enrichment p-value ≤ 1 e-12 were considered significant.

**Supplementary Table 3:** Enrichment of G-quadruplex forming motifs in ORC1 ChIP data. G4Hunter was used (with a stringent threshold of 1.7 to identify high-confidence motifs) to identify G4 forming motifs in ORC1 ChIP peak and summit sequences at 24 and 30 hpi, and in the whole genome. Enrichment was calculated as the ratio of the total number of G4 per bp in the peak or summit sequences and the total number of G4 per bp in the whole genome.

|  |  | (n) sequences | total length | G4 count | Enrichment |
| --- | --- | --- | --- | --- | --- |
| **ORC1 ChIP peaks** | 24 hpi | 8843 | 4599930 | 101 | 2.05 |
|  | 30 hpi | 6785 | 3337581 | 101 | 2.82 |
| **ORC1 ChIP summits**<br>**± 50 bp** | 24 hpi | 12420 | 1242000 | 8 | 0.60 |
|  | 30 hpi | 8997 | 899700 | 13 | 1.35 |
| **whole genome** |  | 16 | 23332831 | 249 |  |

**Supplementary Table 4:** Spearman correlation between ORC1 ChIP data and known epigenetic markers. Genome-wide correlation coefficients between ORC1 ChIP log2ratios at 24 and 30 hpi and log2ratios of histone markers, chromatin accessibility, as well as HP1 placement at different timepoints in the parasite lifecycle were calculated using deeptools multiBigwigSummary (bin size of 100 bp). Correlation coefficients between ORC1 ChIP and HP1 outside telomeric regions were also calculated. Negative correlations are highlighted in red and positive correlations in blue.

|  |  | ORC1 ChIP |  |
| --- | --- | --- | --- |
|  |  | 24 hpi | 30 hpi |
| AP2 |  | -0.2078 | -0.1918 |
| ATACseq | 5 hpi | -0.3085 | -0.2965 |
|  | 10 hpi | -0.5751 | -0.5653 |
|  | 15 hpi | -0.4839 | -0.4734 |
|  | 20 hpi | -0.6302 | -0.6138 |
|  | 25 hpi | -0.5514 | -0.5343 |
|  | 30 hpi | -0.6207 | -0.5990 |
|  | 35 hpi | -0.4919 | -0.4714 |
|  | 40 hpi | -0.5764 | -0.5537 |
| H2A | 40 hpi | 0.0338 | 0.0293 |
| H2A.z | 10 hpi | -0.0309 | -0.0335 |
|  | 20 hpi | 0.0069 | 0.0021 |
|  | 30 hpi | 0.0004 | -0.0066 |
|  | 40 hpi | -0.1139 | -0.1141 |
| H3K4me3 | 10 hpi | 0.0009 | -0.0005 |
|  | 20 hpi | 0.0093 | 0.0049 |
|  | 30 hpi | -0.0001 | -0.0030 |
|  | 40 hpi | -0.0331 | -0.0314 |
| H3K9ac | 10 hpi | -0.0007 | -0.0031 |
|  | 20 hpi | 0.0016 | -0.0035 |
|  | 30 hpi | -0.0780 | -0.0812 |
|  | 40 hpi | -0.1493 | -0.1528 |
| HP1 | rings | 0.4434 | 0.4663 |
|  | trophs | 0.3402 | 0.3699 |
|  | schizonts | 0.5485 | 0.5737 |
| HP1<br>(no telomeres) | rings | 0.3764 | 0.3836 |
|  | trophs | 0.2555 | 0.2704 |
|  | schizonts | 0.4993 | 0.5095 |

**Supplementary Table 5:** Spearman correlation between DNAscent active origins and known epigenetic markers. Genome-wide correlation coefficients between DNAscent origin log2ratios (at 30 hpi, 36 hpi and both time points combined; over the whole genome or only at *var* genes) and log2ratios of histone markers, chromatin accessibility, HP1 placement, and gene expression (‘RNA-seq’) at different timepoints in the parasite lifecycle were calculated using deeptools multiBigwigSummary (bin size of 100 bp). Negative correlations are highlighted in red and positive correlations in blue.

|  |  | DNAscent origins log2ratio |  |  |
| --- | --- | --- | --- | --- |
|  |  | 30 hpi | 36 hpi | All |
| DNAscent origins log2ratio | 30 | 1.0000 | 0.0435 | 0.4243 |
|  | 36 | 0.0435 | 1.0000 | 0.9033 |
|  | All | 0.4243 | 0.9033 | 1.0000 |
| ORC1 ChIP log2ratio (whole genome) | 24 | -0.0014 | -0.0534 | -0.0597 |
|  | 30 | -0.0035 | -0.0699 | -0.0733 |
| ORC1 ChIP log2ratio (var genes) | 24 | 0.1298 | 0.2403 | 0.2461 |
|  | 30 | 0.1501 | 0.2933 | 0.3065 |
| AP2 | AP2 | -0.0138 | -0.0048 | -0.0099 |
| ATACseq R1 | 5 | 0.0139 | 0.0088 | 0.0111 |
|  | 10 | 0.0037 | 0.0062 | 0.0066 |
|  | 15 | 0.0064 | -0.0027 | -0.0015 |
|  | 20 | -0.0020 | -0.0027 | -0.0027 |
|  | 25 | -0.0134 | -0.0197 | -0.0217 |
|  | 30 | -0.0200 | -0.0415 | -0.0413 |
|  | 35 | -0.0207 | -0.0628 | -0.0617 |
|  | 40 | -0.0029 | -0.0285 | -0.0267 |
| ATACseq R2 | 5 | 0.0086 | -0.0043 | -0.0025 |
|  | 10 | 0.0141 | 0.0067 | 0.0092 |
|  | 15 | 0.0030 | -0.0005 | 0.0005 |
|  | 20 | -0.0052 | -0.0054 | -0.0070 |
|  | 25 | -0.0213 | -0.0345 | -0.0375 |
|  | 30 | -0.0304 | -0.0557 | -0.0576 |
|  | 35 | -0.0263 | -0.0715 | -0.0714 |
|  | 40 | -0.0049 | -0.0431 | -0.0407 |
|  |  | 30 hpi | 36 hpi | All |
| H2A | 40 | -0.0002 | -0.0025 | -0.0027 |
| H2A.z | 10 | 0.0069 | 0.0111 | 0.0135 |
|  | 20 | 0.0041 | -0.0014 | 0.0005 |
|  | 30 | 0.0160 | 0.0134 | 0.0186 |
|  | 40 | 0.0185 | 0.0449 | 0.0490 |
| H3K4me3 | 10 | 0.0039 | -0.0072 | -0.0034 |
|  | 20 | 0.0012 | -0.0059 | -0.0044 |
|  | 30 | 0.0003 | 0.0002 | 0.0025 |
|  | 40 | -0.0030 | -0.0020 | -0.0001 |
| H3K9ac | 10 | 0.0039 | -0.0040 | -0.0012 |
|  | 20 | 0.0000 | -0.0114 | -0.0098 |
|  | 30 | 0.0138 | 0.0084 | 0.0139 |
|  | 40 | 0.0197 | 0.0121 | 0.0207 |
| HP1 | rings | 0.0074 | 0.0234 | 0.0226 |
|  | schizonts | 0.0014 | 0.0158 | 0.0130 |
|  | trophs | 0.0129 | 0.0288 | 0.0312 |
| RNAseq | 5 | -0.0423 | -0.0773 | -0.0923 |
|  | 10 | -0.0614 | -0.0857 | -0.1046 |
|  | 15 | -0.0747 | -0.0949 | -0.1145 |
|  | 20 | -0.0815 | -0.1118 | -0.1297 |
|  | 25 | -0.0797 | -0.1324 | -0.1459 |
|  | 30 | -0.0734 | -0.1512 | -0.1622 |
|  | 35 | -0.0577 | -0.1662 | -0.1736 |
|  | 40 | -0.0284 | -0.1376 | -0.1420 |

**Supplementary Table 6: Gene Ontology Terms**

List of Gene Ontology (GO) terms of genes that overlap with active DNAscent origins ranked according to total number of occurrences.

**Supplementary Table 7:**
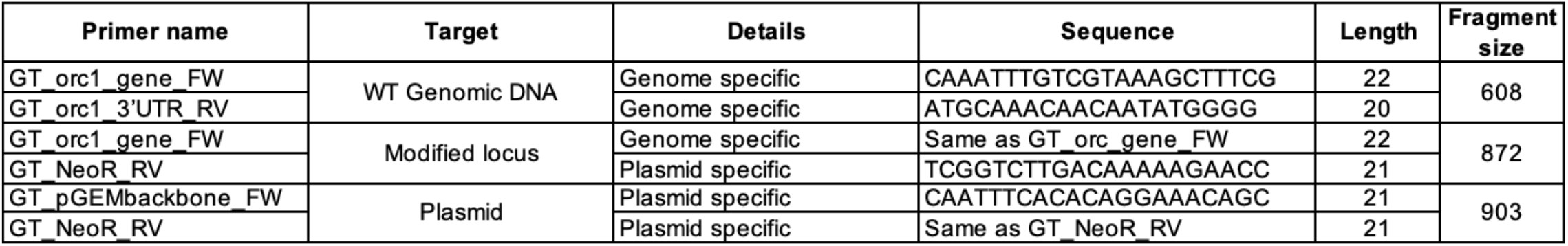
Genotyping primers. Genotyping and control primer sequences used to confirm successful integration of 3xHA and neomycin resistance gene into *orc1* (PF3D7_1203000) and expected fragment sizes.

## Notes

### Competing Interest Statement

The authors have declared no competing interest.

